# The transcription factor Shox2 shapes thalamocortical neuron firing and synaptic properties

**DOI:** 10.1101/660662

**Authors:** Diankun Yu, Matthieu Maroteaux, Yingnan Song, Xiao Han, Isabella Febbo, Claire Namboodri, Cheng Sun, Wenduo Ye, Emily Meyer, Stuart Rowe, YP Chen, LA Schrader

**Author notes:** Address for LAS: Cell and Molecular Biology, 2000 Percival Stern Hall, 6400 Freret St., New Orleans, LA 70118.

## Abstract

Thalamocortical neurons (TCNs) transmit information about sensory stimuli from the thalamus to the cortex. In response to different physiological states and demands TCNs can fire in tonic and/or phasic burst modes. These firing properties of TCNs are supported by precisely timed inhibitory synaptic inputs from the thalamic reticular nucleus and intrinsic currents, including T-type Ca^2+^ and HCN currents. These intrinsic currents are mediated by Cav3.1 and HCN channel subunits, and alterations in expression or modulation of these channels can have dramatic implications on thalamus function. The factors that regulate these currents controlling the firing patterns important for integration of the sensory stimuli and the consequences resulting from the disruption of these firing patterns are not well understood. Shox2 is a transcription factor known to be important for pacemaker activity in the heart. We show here that Shox2 is also expressed in adult mouse thalamus. We hypothesized that genes regulated by Shox2’s transcriptional activity may be important for physiological properties of TCNs. In this study, we used RNA sequencing on control and *Shox2* knockout mice to determine *Shox2*-affected genes and revealed a network of ion channel genes important for neuronal firing properties. Quantitative PCR confirmed that expression of *Hcn*2, *4* and *Cav3*.*1* genes were affected by *Shox2* KO. Western blotting showed expression of the proteins for these channels was decreased in the thalamus, and electrophysiological recordings showed that Shox2 KO impacted the firing and synaptic properties of TCNs. Finally, behavioral studies revealed that *Shox2* expression in TCNs play a role in somatosensory function and object recognition memory. Overall, these results reveal Shox2 as a transcription factor important for TCN firing properties and thalamic function.

## INTRODUCTION

Processing of sensory information is mediated by precise circuitry that senses stimuli in the periphery and transforms the information through a network of synaptic connections to ultimately allow perception and cognitive processing of the surrounding world. Rhythmic oscillations of brain activity crucial to cognitive function emerge from neuronal network interactions that consist of reciprocal connections between the thalamus, the inhibitory thalamic reticular nucleus and the cortex^1^. Dysfunction of these oscillations caused by aberrant activity in the thalamic circuit is thought to play a role in many neuropathological conditions, including epilepsy ^2-4^, autism ^5-7^, and schizophrenia ^8-11^. Furthermore, damage to thalamic nuclei, especially medial and anterior nuclei, causes severe memory deficits known as diencephalic amnesia ^12-17^.

The intrinsic properties that shape action potential firing and contribute to rhythmic oscillations of the thalamocortical neurons (TCNs) are important for efficient transfer of information from the thalamus to the cortex. Notably, TCNs switch their firing states between 2 modes, burst and tonic firing modes that occur at different membrane potentials ^18-20^. The transitions between tonic and burst modes are controlled by several intrinsic conductances within the TCNs, mainly T-type Ca^2+^ currents (I_T_) and H-currents (I_h_), mediated by Cav3.x and HCN family of channels, respectively ^1^. Interplay of the kinetics of these currents and other conductances, including leak K^+^, inwardly rectifying K+, a persistent Na current and transient A-type currents, can establish a cycle of oscillatory activity that generates rhythmic activity in the thalamocortical network ^21^. Consequently, modulation of the intrinsic properties of TCNs is an important regulatory mechanism of firing activity crucial for sensory perception, sleep activity and cognition. Very little is understood about the factors that contribute to modulation of the ion channels that underlie these currents.

The transcription factor, Shox2, represents a possible mechanism to coordinate expression of ion channels important for TCN firing properties. Shox2 is a member of the homeobox family of transcription factors ^22-24^, and recent studies suggest it is important for development and maintenance of rhythmic activity in adult heart. Cells in the sinoatrial node of the heart generate spontaneous, rhythmic action potentials and lead other working myocytes to beat at a stable firing rate, thus these cells are known as pacemaker cells ^25^. The rhythmic action potential generation in the pacemaker cells is mediated by the prominent expression of HCN channels and T-type calcium channels ^26, 27^. We and others have shown that *Shox2* plays a decisive role in the differentiation of pacemaker cells in the sinoatrial node of the heart and pulmonary vein in mice, and Shox2 expression in the SAN is necessary for cardiac pacemaker-type activity ^28, 29^. Importantly, Shox2 is essential for expression of HCN4 channels ^28^, and *Shox2* overexpression favors differentiation of mouse embryonic stem cells into pacemaker cells with biological pacing ability and enhanced HCN currents^30^. *Shox2* expression in the sinoatrial node of the heart continues into adulthood since cells of the sinoatrial node maintain pacemaker properties, while *Shox2* expression is suppressed in pulmonary vein and coronary sinus in the adult, and these cells lose pacemaker properties during development^31^. These results suggest that Shox2 expression is critical for expression of channels important for myocyte firing properties and is a determinant of pacemaker properties of the sinoatrial node.

During development of the nervous system, *Shox2* expression has been reported in neurons of the facial motor nucleus (nVII) ^32^, cerebellum ^33^, spinal cord ^34^ and dorsal root ganglia ^35^. *Shox2*-expressing excitatory interneurons in the ventral spinal cord are rhythmically active during locomotor-like activity, and synaptic and electrical connections between Shox2-expressing interneurons are crucial for the frequency and stability of their rhythmic activity ^34, 36^, showing that interconnectivity between *Shox2*-expressing neurons is critical for synchronization of rhythmic activity. These results suggest that *Shox2*-expressing neurons play a critical role in central pattern generation important for locomotion, but the role of *Shox2* in this rhythmic behavior was not further investigated.

In this study, we found that *Shox2* is expressed in thalamocortical neurons (TCNs) in adult mice. TCNs express HCN2, HCN4 and Ca_v_3.1 channel protein subunits that are important for firing properties of TCNs, and we hypothesized that *Shox2* coordinates the expression of genes for these ion channels to affect action potential firing activity of TCNs. Using conditional KO animals, we further demonstrated that *Shox2* is important for tonic spike frequency firing properties in TCNs, likely by affecting expression multiple ion channels, including HCN. We also demonstrated that an inducible global knock-out of *Shox2* reduced anxiety behavior in the open field, impaired sensorimotor function and object recognition memory. Object recognition memory deficits were confirmed with an inducible Shox2 knock-out in medial thalamus. The present study provides novel insight into the molecular markers that contribute to thalamocortical neuron function and shows that S*hox2* expression is critical to maintain thalamic neuron function and physiological properties by regulating gene expression in the neurons of the adult mouse thalamus.

## Methods

### Mice

All animal procedures were approved by Tulane University Institutional Animal Care and Use Committee (IACUC) according to National Institutes of Health (NIH) guidelines. *Shox2* transgenic mice including *Shox2*^*Cre*^, *Shox2*^*LacZ*^, *Shox2*^*f/f*^ and *Rosa26*^*CreERt*^ mice were generously donated by Dr. Yiping Chen. All wildtype C57Bl/6N mice were ordered from Charles River. *Rosa26*^*LacZ/+*^ *(stock #003474) and Gbx2*^*CreERt/+*^ breeders (stock #022135) were ordered from Jackson Lab.

In inducible KO experiments, *Rosa26*^*CreERt/+*^,*Shox2*^*f/f*^ or *Rosa26*^*CreERt/CreERt*^,*Shox2*^*f/f*^ female mice were crossed with *Shox2*^*-/+*^ male mice or, in the case of the *Gbx2* animals, *Gbx2*^*CreERt/+*^,*Shox*^*f/f*^ or *Gbx2*^*CreErt/CreErt*^,*Shox*^*f/f*^ were crossed with *Shox2*^-/+^ male mice (Supplemental Fig. 1). Litters were labelled and genotyped at postnatal day 10 (P10). The KO group was the *Rosa26*^*CreERt/+*^, *Shox2*^*-/f*^ mice (*Gbx2*^*CreRt/+*^, *Shox2*^*-/f*^) and the control (CR) group was the littermate *Rosa26*^*CreERt/+*^ *(Gbx2*^*CreErt/+*^*), Shox2*^*+/f*^. In the *Shox2*^*LacZ/+*^ and *Shox2*^*Cre/+*^ mice, the first two exons of the *Shox2* allele were partially replaced by *LacZ* and *Cre* genes respectively in order to obtain the expression of *LacZ* mRNA and *Cre* mRNA under the control of the *Shox2* promoter, while the unaffected alleles express *Shox2* mRNA ^24^. The *Rosa26*^*CreERt/+*^ mouse line is a transgenic mouse line with a tamoxifen-inducible Cre^ERt^ inserted in the *Rosa26* loci. The *Rosa26*^*LacZ/+*^ mouse line is a transgenic mouse line with a floxed stop signals followed by LacZ gene inserted in the *Rosa26* loci ^37^. This ‘cre reporter’ mouse strain was used to test the expression of the *Cre* transgene under the regulation of a specific promoter.

The Rosa26^CreErt^ is a global KO, whereas the *Gbx2*^*CreERt*^ mouse was used to knockdown *Shox2* specifically in the medial thalamus. Our localization studies with *Gbx2*-promotor driven GFP staining showed that *Gbx2* promotor-driven *CreERt* is specifically expressed in the medial thalamus of the Gbx2^CreErt^ adult mouse (Supplemental Fig. 2A). Further testing using RT-qPCR showed that *Shox2* mRNA was reduced in the medial thalamus of the *Gbx2*^*CreRt/+*^, *Shox2*^*-/f*^ compared to CR mice, but not lateral thalamus (Supplemental Fig. 2B).

To induce recombination in animals bearing a Cre^ERT2^, pre-warmed tamoxifen (100-160 mg/kg) was injected intraperitoneally into KO mice and CR littermates at the same time every day for five consecutive days. Tamoxifen (20 mg/mL) was dissolved in sterile corn oil (Sigma, C8267) with 10% alcohol. The littermate KO mice and CR mice of the same sex were housed together and received the same handling. Throughout experiments, the researchers were blinded to the genotype. RT-qPCR experiments were used to confirm the efficiency of *Shox2* KO in brains in every animal tested.

In order to view projections of *Shox2*-expressing neurons, we created the Ai27D-*Shox2Cre* mouse. B6.Cg-Gt(ROSA)26Sortm27.1(CAG-COP4*H134R/tdTomato)Hze/J(Ai27D) mice from Jackson labs were crossed with *Shox2*Cre to obtain mice with Shox2-expressing neurons labeled with tdTomato and expressing ChR2 (figure 2 M-O).

### X-gal staining

Adult *Shox2*^*LacZ/+*^ or *Shox2*^*Cre/+*^, *Rosa26*^*LacZ/+*^ male mice were anaesthetized by isoflurane inhalation, decapitated, and the brains were removed. Brains were sliced at 200 μm using a Vibratome Series 3000 Plus Tissue Sectioning system. Brain slices were placed into ice-cold artificial cerebrospinal fluid (aCSF) in a 24-well plate and fixed with 0.5% glutaraldehyde and 4% paraformaldehyde in phosphate buffered saline (PBS) for 15 minutes. After 3X wash with ice-cold PBS, the slices were incubated with X-gal staining solution, containing: (X-gal (1mg/ml), potassium ferrocyanide (4 mM), potassium ferrcyanide (4mM) and MgCl_2_(2 mM) and covered by aluminum foil at 37 °C overnight. All slices were washed and post-fixed. Images were taken under a stereo microscope.

### Immunohistochemistry (IHC)

Mice were deeply anaesthetized by injection with ketamine (80 mg/kg) mixed with xylazine (10 mg/kg), perfused transcardially with ice-cold phosphate-buffered saline (PBS) followed by 4% paraformaldehyde in PBS and decapitated for brain collection. Mouse brains were placed in 4% paraformaldehyde in PBS at 4° C overnight for post-fixation. In order to perform cryostat sections, the brains were sequentially placed in 15% and 30% sucrose in PBS solutions at 4 °C until saturation. The brain samples were embedded with optimal cutting temperature compound (OCT) and stored at −20 °C and cryo-sectioned in 20-50 μm coronal slices with Leica CM3050S cryostat. For IF staining, slices were washed with 50 mM Tris Buffered Saline with 0.025% Triton X-100 (TTBS) and blocked in 2% Bovine Serum Albumin (BSA) in TTBS for 2 hours at room temperature. Primary antibodies were diluted in blocking solutions and applied on slides overnight at 4°C. Fluorescence-conjugated secondary antibodies were diluted 1:1000 in blocking solutions and applied on slides for one hour at room temperature. 1:1000 DAPI was applied for 5 minutes at room temperature for nuclei staining and washed. The slices were mounted on slides with mounting media (Vector Laboratories, H-1000) and imaged under confocal microscope.

### mRNA Sequencing

Thalamus mRNA was extracted from 3 KO mice and 3 CR mice and sent to *BGI Americas Corporation (Cambridge, MA, USA)* for RNA-seq quantification. Total RNA was isolated in tissue from the midline of the thalamus of P60 male *Gbx2*^CreERt/+^, *Shox2*^-/f^ mice and control male littermates (*Gbx2*^CreERt/+^, *Shox2*^+/f^) with the same method used for RT-qPCR RNA extraction. Around 30 million single-end 50-bp reads by BGISEQ-500 Sequencing Platform per sample were aligned to the mm10/GRCm38 mouse reference genome using Salmon v0.10.2 ^38^. The count data from Salmon v0.10.2 was analyzed via DESeq2 v 1.22.2 ^39^ to identify genes differently expressed (DEGs) between KO and CR and to calculate Fragments Per Kilobase Million (FPKMs). Genes with adjusted p value < 0.1 were defined as DEGs and were used for further gene ontology (GO) analysis through online DAVID Bioinformatics Resources ^40, 41^.

### Quantitative reverse transcription PCR (RT-qPCR)

The whole thalamus was collected from adult mouse brains *(Rosa*^*CreErt-*^*Shox2* KO) and immediately stored in RNAlater™ RNA stabilization reagent (ThermoFisher Scientific, AM7021). RNA was extracted using RNeasy Mini Kit (Qiagen, 74104) following the standard protocol provided in the manual. The RNA concentration and quality were tested using Nanodrop Microvolume Spectrophotometers and Fluorometer as well as agarose gel investigation. Reverse transcription was conducted using iScript™ Reverse Transcription Supermix (Bio-Rad, 1708840). Quantitative PCR was conducted with iTaq™ Universal SYBR Green Supermix (Bio-rad, 1725121) in Bio-Rad CFX96 Touch™ PCR system. Data analysis was done with CFX Manager software. The expression of all the genes tested in the RT-qPCR experiments were normalized to the widely used housekeeping reference gene β-actin (*Actb*) and TATA-box binding protein (*Tbp*) ^42, 43^. All primers were designed and tested, and conditions were optimized to have an efficiency between 95% and 105%. Both the melt curves and gel investigations were used to confirm the RT-qPCR products. The primer sequences of all tested genes including reference gene *Actb* and *Tbp* are listed in Table 1.

**Table 1.**
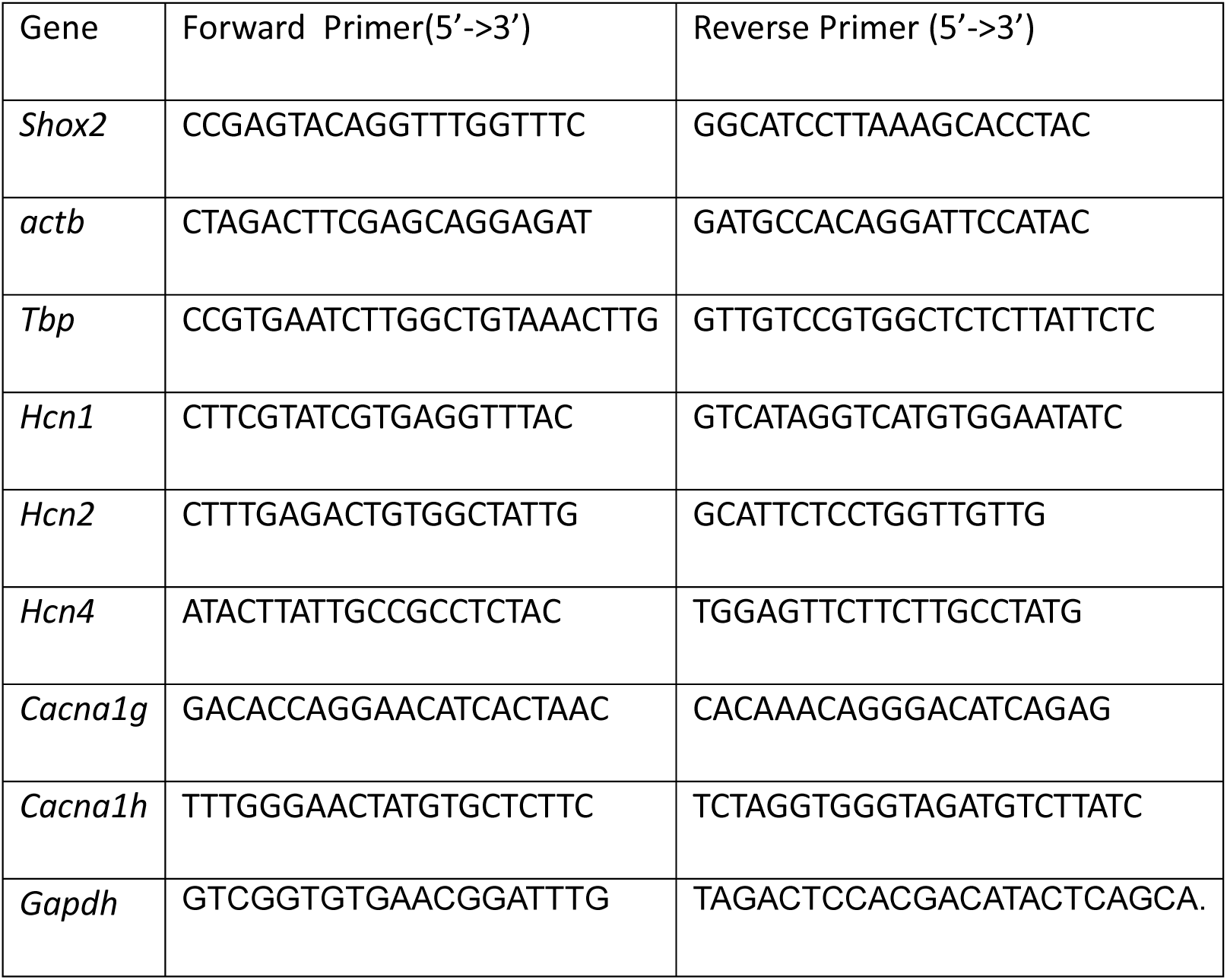
Sequences of RT-qPCR primers.

To confirm knock-down of *Shox2* in the Gbx2^CreERt/+^ animals, adult *Gbx2*^*CreERt/+*^; *Shox2* ^*-/f*^ male and female mice were anaesthetized by isoflurane inhalation followed decapitation. The brains were removed and a 1 mm thick slice through the thalamus was removed via razor blade, the location of cut is determined by Paxinos and Franklin Mouse Brain Atlas. Medial thalamus tissue is collected with 1mm stainless steel punching tool and lateral thalamus was separated via razor blade. The collected tissues were stored in 50 μL RNA later solution and stored in −80 freezer. To homogenize collected tissues, 350 μL of RLT lysis buffer from Qiagen RNeasy Mini Kit is added to the tissue and homogenized with a pestle mortar. The homogenized tissues went through sonication with a Q55 sonicator (Qsonica) and then 350 μL cold, 70% EtoH was added to the sample. After this, the mixed solution is processed by series of spin and wash follow the instructions book from Qiagen RNeasy Mini Kit. Once RNAs are isolated from tissues, we applied qRT-PCR with Shox2 primer (Table 1) and normalized with GAPDH.

### Western Blot

Thalamic tissues were collected from the adult mouse brains and immediately placed on dry ice and stored at −80 °C until use. The thalamus samples were lysed with RIPA lysis buffer (150 mM Sodium chloride, 1% Triton X-100, 0.5% sodium deoxycholate, 0.1% SDS, 50 mM Tris, pH 8.0) with fresh added Halt™ protease inhibitor cocktail (ThermoFisher Scientific, 78430). Samples were centrifuged at 12,000 rpm at 4 °C for 20 minutes and the supernatant protein samples were collected. The protein concentration of the samples was determined using the Bio-Rad DC protein assay (Bio-Rad, 500-0116). Samples were normalized with the same lysis buffer, aliquoted and stored at −80°C until use. Before loading, 5X sample buffer (ThermoFisher Scientific, 39001) and dithiothreitol (final concentration - 50 mM, DTT) were added to each protein sample. Sample mixtures were left at room temperature for 30 minutes. Protein (20-30 μg/well) was loaded in a SDS-PAGE gel (4% stacking gel and 8% separating gel), together with 3 μL prestained protein ladder (ThermoFisher Scientific, 26619). The gels were run at 70 mV for 3 hours, and the proteins were transferred to a pre-activated PVDF membrane (Millipore, IPFL00005) at −70 mV for 3 to 4 hours. Sodium dodecyl sulfate (SDS) and methanol were added into transfer buffer at a final concentration of 0.1% and 10%, respectively. The gels were stained with Coomassie Brilliant Blue solution (0.1% Coomassie Brilliant Blue, 50% methanol, 10% Glacial acetic acid) to check that no obvious proteins remained under these transfer conditions. Membranes were incubated in blocking solution with 5% non-fat dry milk and 3% BSA in TTBS at room temperature for one hour. Primary antibody was diluted in Odyssey^R^ Blocking Buffer in TBS and applied on the membrane at 4 °C overnight. After washing the membrane with TTBS, fluorescence-conjugated secondary antibodies were diluted and applied on the membrane at room temperature for one hour. Imaging of the stained membrane was done in an Odyssey CLx Infrared Scanner and analyzed by Image Studio Lite Ver 5.2.

### Antibodies used in IHC and Western blot experiment

| Reagent/Resource | Supplier | Details |
| --- | --- | --- |
| Chicken anti-β-galactosidase | Abcam | Ab9361; 1:500 |
| Rabbit anti-NeuN Cy3- conjugated | Millipore Sigma | ABN78; 1:500 |
| Mouse anti-parvalbumin | Millipore Sigma | MAB1572; 1:500 |
| Rabbit-anti GFAP | PhosphoSolutions | 620-GFAP; 1:300 |
| Rabbit-anti GFP | Novus Biologicals | NB 600-308; 1:300 |
| Alexa-Fluor 488 Goat anti-mouse | Life Technologies | 1:1000 |
| Alexa-Fluor 647 Goat anti-chicken | Life Technologies | 1:1000 |
| Alexa-Fluor 594 Goat anti-rabbit | Life Technologies | 1:1000 |
| mouse anti-HCN2 | Neuromab | N71/37; 1:1000 |
| mouse anti-HCN4 | Neuromab | N114/10; 1:1000 |
| mouse anti-Cav3.1 | Neuromab | N178A/9; 1:1000 |
| IRDye 680RD Goat anti-mouse | Li-Cor | P/N 926-68070; 1:10,000 |
| IRDye 800CW Goat anti-rabbit IgG (H+L) | Li-Cor | P/N 926-32211; 1:10,000 |

### Electrophysiology

At the same time of the day (11:00 am summer and 10:00 am winter), male and female mice (PND 60-120) were anaesthetized with isoflurane and decapitated. Brains were quickly removed and immersed in oxygenated (95% O_2_ and 5% CO_2_), ice-cold N-methyl-D-glucamine (NMDG)-based slicing solution (in mM, 110 NMDG, 110 HCl, 3 KCl, 1.1 NaH_2_PO_4_, 25 NaHCO_3_, 25 Glucose, 10 ascorbic acid, 3 pyruvic acid, 10 MgSO_4_, 0.5 CaCl_2_). The first 350 μM coronal brain section containing the most anterior paraventricular thalamus (PVA) was obtained with a Vibratome Series 3000 Plus Tissue Sectioning System. The collected brain slices were transferred and incubated in oxygenated standard aCSF (in mM, 125 NaCl, 2.5 KCl, 26 NaHCO_3_, 1.24 NaH_2_PO_4_, 25 Dextrose, 2 MgSO_4_, 2 CaCl_2_) at 37 °C for 30 minutes, then incubated at room temperature until use.

During the recordings, an individual slice was transferred to a recording chamber and perfused with oxygenated external solution at a speed of 1 mL/minute at room temperature at room temperature. Unless otherwise specified, standard aCSF was perfused as the external solution. For isolation of the specific currents, different pharmacological antagonists were applied in the external solution as stated in the results. PVA was identified as the nucleus near the border of the third ventricle enclosed by the stria medullaris. Cell-attached and whole-cell recordings were obtained using MultiClamp 700B amplifier, Digidata 1322A digitizer, and a PC running Clampex 10.3 software (Molecular Device). For cell-attached recording, glass pipettes had resistances of 2.5 – 3.5 MΩ filled with standard aCSF. Giga seals were obtained in every cell by application of a small negative pressure for spontaneous action potential recording. For intracellular whole-cell patch clamp recording, glass pipettes had resistances of 3.5 – 6 MΩ filled with internal pipette solution (in mM, 120 Kgluconate, 20 KCl, 0.2 EGTA, 10 Hepes, 4 NaCl, 4 Mg^2+^ATP, 14 phosphocreatine, 0.3 Tris GTP (pH was adjusted to 7.2-7.25 by KOH, osmolarity was adjusted to 305-315 mOsm by sorbitol). Series resistance was monitored and only cells with series resistance less than 20 MΩ and that did not change over 15% throughout the recording were further analyzed. Spontaneous action potentials were recorded in current clamp mode at membrane potential. Cells with no action potentials identified in 5 minutes are classified as ‘not active’ cells. Action potential threshold was measured on the first spike at the point where the voltage change reaches 20 mV/ms.

HCN currents were isolated under voltage-clamp. The external solution for HCN current isolation contained 0.5 μM TTX, 1 mM NiCl_2_, 1 mM CdCl_2_, 2 mM BaCl_2_, 10 μM DNQX and APV to block voltage-gated sodium channels, voltage-gated calcium channels, inwardly-rectifying potassium channels and excitatory synaptic current respectively. NaH_2_PO_4_ was omitted to prevent precipitation with cations.

### Behavioral assays

If a timeline is needed, P21-P25 tamoxifen injection; ∼around P56 1-2 weeks handling; then 3 day habituation of the room, 1 day open field, 2 days NOR, 1 weeks later paw sensation, 1 weeks later forced swim, at least 1 weeks later tail suspension. 3 days handling before each test.

An open field test was used to test mouse exploratory behavior and anxiety-related behavior. The experiments were all done 2 hours into the animal’s dark light phase under dim red light. All mice received routine handling for a week. Three days before the experiments, mice were habituated to the training and testing room for 1 hour each day. On the first day of experiments, each mouse was placed in the open field (16 inches x 16 inches) and allowed to explore freely under dim red light for 5 minutes. Infrared beams and computer-based software *Fusion* were used to track mice and calculate mice activity and time spent in the center (8 inches x 8 inches) of total open field.

In novel object recognition (NOR) experiments, all mice received routine handling and three days habituation to the experimental room before the experiments. On the day before the familiarization trial, each mouse was placed in the open field in the absence of objects and allowed to explore it freely, the behaviors were recorded and analyzed further as open field test data. In the familiarization trial, each mouse was placed in the open field containing two identical 100 ml beakers in the neighboring corners for 5 minutes. Twenty-four hours later, each mouse was placed back in the same open field with two objects, one of which was the 100 ml beaker and the other one a padlock of a similar dimension, for a 5-minute testing trial. To prevent bias in objects exploration, mice were always released on the opposite side from the object for both familiarization and testing trials. For NOR and the subsequent behavior experiments, mouse behaviors in the testing trials were video-taped and analyzed by experimenters who were blind to the genotypes of the mice. Exploration of an object was defined as sniffing and touching the object with attention, whereas other behaviors like running around the object, sitting or climbing on it were not recorded as exploration ^48^. Discrimination index was calculated as (t_n_ - t_f_)/(t_n_ +t_f_). Where t_n_ = time exploring new object and t_f_ = time exploring familiar object. In the *Gbx2Cre,Shox2* KO test, 2 animals exhibited a preference for one object in the familiarization trial and therefore were not tested the next day.

The adhesive removal test was used to assess mouse paw sensorimotor response ^49^. A small piece of round sticky paper tape (Tough-spots, for microtube cap ID, ∼1 cm^2^, Research Products International Corp. 247129Y) was applied to the plantar surface of the right hind paw of each mouse, and the mouse was placed back in its home cage and the behavior recorded. The latency to the first response to the tape was measured and analyzed.

The tail suspension test and forced swim test were applied to assess and evaluate mouse depressive-like behaviors ^50^. In the tail suspension test, a 5-cm of the tip of Falcon 15 mL conical centrifuge tube was placed around the tails to prevent tail climbing, and each mouse was suspended by the tail for 5 minutes. The behavior of the mice was recorded and analyzed. The escape-related struggling time within the 5-minute experiment was measured as mobility time.

In the forced swim test, each mouse was placed in a 1000 ml beaker with ∼800 ml water for 5 minutes. The behavior of the mouse was taped and analyzed by experimenters who were blinded to genotypes of the mice. Swimming and intentional movements with all four legs or body were measured as mobility time, and small movements of front or hind legs made by the animal to stay at the surface were not counted as mobility.

Fear Conditioning: Fear conditioning (habituation, training and testing) was performed and filmed in standard operant chambers (Med Associates, video fear conditioning). All behaviors were recorded, and mobility or freezing behavior was assessed online by Medical Associates Video Freeze software.

The fear conditioning protocol occurred over 4 days as follows:

Day 1: Habituation - each animal was placed in the operant chamber for 10 minutes.

Day 2: Training - each animal was placed in the chamber for a total of 8 minutes. The training trial was two mild training sessions consisting of a 30 sec auditory cue (administered at 3 and 5 minutes after placement in chamber) that co-terminated with a single 2 second 0.5 mA shock. The animal was removed from the chamber after 8 minutes.

Day 3: Context Testing: Animal was placed in the chamber for 5 minutes and behaviors recorded.

Day 4: Cued testing - the chamber was modified (plastic floor and inserts to allow different shape), and different olfactory cues (vanilla) were given. The cue was administered after 3 minutes chamber exploration and freezing to the cue was assessed as described above. Animals were removed from the chamber after 6 minutes.

### Statistics

Unless otherwise noted in the text, control and KO results were compared using a Student’s unpaired t-test. In some cases of unequal variance, the Mann Whitney nonparametric test is used instead of t-test.

## Results

### *Shox2* is specifically expressed in the thalamic neurons in the brain of young adult mouse

To investigate the expression of *Shox2* in postnatal mouse brain, coronal brain slices from P25 and P60 *Shox2*^*LacZ/+*^ mice were stained with X-gal, in which the expression of *LacZ* indicated *Shox2* expression (Supplemental Fig. 3 A, B). The X-gal staining results indicated that *Shox2* is expressed throughout the dorsal thalamus including medial thalamus, anterior thalamus nuclei (ATN), ventrobasal nucleus (VB), dorsal lateral geniculate nucleus (dLGN), and medial lateral geniculate nucleus (MGN) but not in other regions of diencephalon including habenula, reticular nucleus of the thalamus and hypothalamus, or other regions of the nervous system like the cortex, subcortical regions of the forebrain, hippocampus, amygdala, cerebellum and spinal cord. To determine whether the expression pattern of *Shox2* changes during development, coronal brain slices from a P56 *Shox2*^*Cre/+*^, *Rosa26*^*LacZ/+*^ mouse in which LacZ is expressed in all cells that have expressed *Shox2* at any time during development were stained with X-gal (Supplemental Fig. 3C). These results showed that the expression of *Shox2* is relatively restricted to the dorsal thalamus in the adult as well as during development, with sparse expression extended to habenula and superior and inferior colliculus and nuclei within the brainstem in the developing animal. We further assessed the cell type of *Shox2*-expressing cells.

In order to determine the cell types in which *Shox2* is expressed, thalamic neurons were labeled with the neuronal nuclear protein antibody (NeuN) which is specifically expressed in mature neurons ^51, 52^. *Shox2* was co-expressed in most, but not all, NeuN-positive cells throughout the dorsal thalamus from rostral to caudal (Fig. 1 A-C). Importantly, all *Shox2*-labeled cells were NeuN-positive, suggesting that *Shox2* expression is restricted to neurons. To confirm that *Shox2* was not expressed in astrocytes, the co-expression of *Shox2* and Glial Fibrillary Acidic Protein (GFAP), which labels astrocytes ^53, 54^ was assessed. GFAP was expressed at relatively low levels in the thalamus but highly expressed in the hippocampus (Supplemental Fig. 4), and *Shox2* was not expressed in the GFAP-positive astrocytes throughout the thalamus (Fig. 1D-F). Together these results show that *Shox2* is expressed in neurons and not GFAP-positive astrocytes.

**Figure 1.**
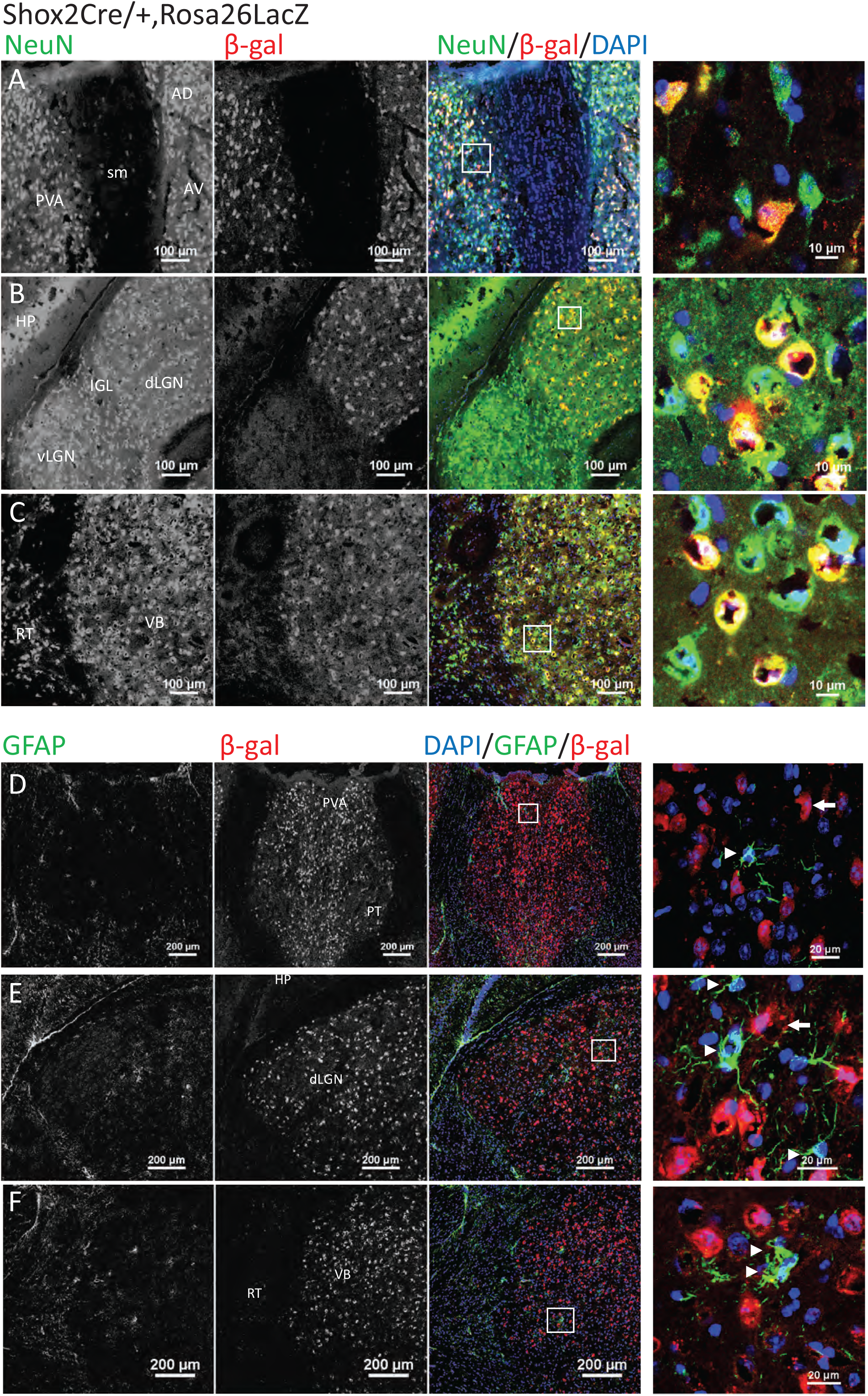
*Shox2* is expressed in NeuN+ neurons in the thalamus and not in GFAP+ astrocytes. Coronal brain sections through the thalamus of *Shox*2^Cre/+^, Rosa26LacZ mice were co-stained with NeuN (green) and β-gal (red). Three typical thalamic regions are shown, including anterior paraventricular thalamus (PVA) (**A**), dorsal lateral geniculate nucleus (dLGN) (**B**), and ventrobasal nucleus (VB) (**C**). Slices were co-stained for NeuN and the reporter for *Shox2*, β-gal. *Shox2* is expressed in NeuN+ neurons (red, merged). Right panels are magnifications of the boxed regions respectively, showing cells that co-express *Shox2* and NeuN. **D-F** show the co-expression of astrocyte marker GFAP (green) and β-gal (red), in three thalamic regions: PVA (**D**), dLGN (**E**) and VB (**F**). Right panels are magnifications of the boxed regions. The arrowheads show the GFAP+ glia, and the white arrows show *Shox2*-expressing cells as indicated by β-gal. No cells co-expressed GFAP and *Shox2*. RT: reticular thalamus. AV: anteroventral nucleus of the thalamus; AD: anterodorsal nucleus of the thalamus. HP: hippocampus.

To determine if *Shox2* is expressed in GABAergic neurons, immunohistochemistry (IHC) with parvalbumin (PV) on coronal brain sections from Shox2^Cre/+^, Rosa26^LacZ/+^ mice was performed (Fig. 2). Parvalbumin (PV) is highly expressed in the interneurons of the reticular nucleus of thalamus, which borders the thalamus laterally and ventrally. PV labeling delineated the reticular nucleus that defined the border of the thalamus. The PV staining results confirmed the results shown in supplemental figure 4 that during development, *Shox2* is expressed throughout the thalamus (Fig. 2) and sparsely in the habenula (Fig. 2D) and midbrain (Fig. 2I). Importantly, *Shox2* was not expressed at any point during development in cells of the cortex (Fig. 2C), reticular nucleus of the thalamus (Fig. 2E, F), hypothalamus (Fig. 2G) or hippocampus (Fig. 2I). In addition, our results showed few PV+ cells in the thalamus as previously reported ^55^, and *Shox2* was not co-expressed in PV+ cells (Fig. 2G). These results suggest that *Shox2* is not expressed in parvalbumin-expressing inhibitory interneurons.

**Figure 2.**
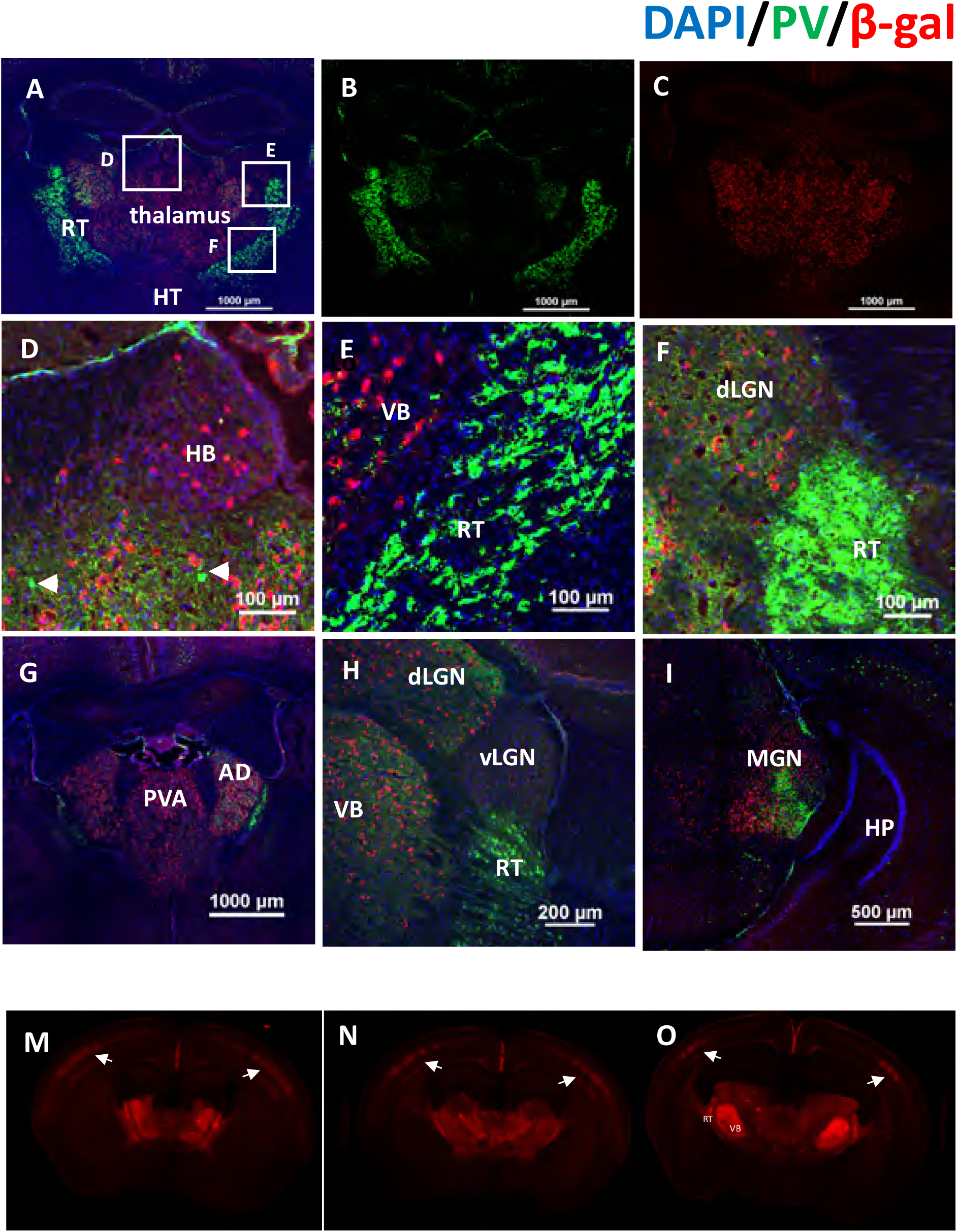
*Shox2* is expressed in glutamatergic thalamocortical neurons but not parvalbumin+ interneurons. Coronal sections through the thalamus were co-stained with parvalbumin (green, **A, B**) and β-gal (red, **A, C**). Boxes in Figure **A** are magnified in **D**. (habenula), **E**. Ventrobasal (VB) and reticular nucleus (RT), and **F**. dorsal lateral geniculate nucleus (dLGN). **G**. Panels G-I show *Shox2* expression in coronal slices of rostral to caudal thalamus (**G**: Paraventricular nucleus of the thalamus (PVA) relative to Bregma, approx. −1.1; **H**: lateral geniculate nucleus (LGN) −2.0; **I**: Medial geniculate nucleus (MGN) –2.9). **M-O:** Coronal sections from rostral (M) to caudal (O) from the Ai27D-*Shox2Cre* in which the presence of tdTomato indicates *Shox2*-expressing neurons. *Shox2* is expressed in neurons throughout the thalamus and projections to the cortex, strongly targeting layer IV barrel cortex (white arrows) and layer VI.

Finally, the expression and projections of *Shox2-*expressing neurons were investigated using *Shox2*^*Cre/+*^; *Rosa*^*tdTomato-ChR2*^ mice. These mice allow labeling of *Shox2*-expressing neurons with td-Tomato and manipulation of *Shox2*-expressing neurons with Channelrhodopsin-2. Interestingly, we found that the *Shox2*-expressing neurons projected to multiple cortical areas, including retrosplenial and somatosensory cortices with a clear delineation of the barrel fields in somatosensory cortex (Fig. 2M-O). Further strong projections from *Shox2*-expressing neurons were observed within the thalamus, particularly the VB complex, in reticular nucleus of thalamus and internal capsule (Fig. 2 M-O) projecting to somatosensory cortex. In summary, the X-gal staining and immunofluorescence results indicated that *Shox2* expression was restricted to excitatory thalamocortical neurons in the adult stage.

### Lack of Shox2 affects gene expression in TCNs

To study the specific role of Shox2 in regulation of gene expression in neurons of adult thalamus, RNA sequencing was performed. For this experiment, the inducible knockout (KO, n = 3) of *Shox2* in *Gbx2*^CreERt/+^, *Shox2*^-/f^ mice were compared to littermate *Gbx2*^CreERt/+^, *Shox2*^+/f^ control (CR, n = 3) mice. Our GFP staining results showed *Gbx2* is specifically expressed in the medial thalamus from P21 (Supplemental fig. 2), so we ran mRNA sequencing with RNA extracted from medial thalamus of the CR and KO mice. Our results showed 372 differentially expressed genes (DEG) between CR and KO mice, 212 of which are downregulated and 160 are upregulated in KO tissue (Supplemental file and Fig. 3A). Gene Ontology (GO) analysis showed *Shox2* KO affected genes in GO terms of ion channel activity, learning and locomotory behavior (Fig. 3B). Importantly, *Shox2* KO downregulated the expression of *Hcn2, Hcn4* and *Cacna1g* genes (Supplemental file and Fig. 3A). The protein products of these genes, HCN2, HCN4 and Cav3.1 mediate HCN current and T-type Ca^2+^ currents, respectively. Since these channels are significant contributors to the rhythmic firing properties of TCNs, we further pursued mRNA and protein expression of these channels.

To confirm *Shox2* regulates ion channel-related genes in the whole thalamus, another transgenic mouse line, the global KO (*Rosa26*^*CreERt/+*^, *Shox2*^*f/-*^ mice), in which *Shox2* was reduced in the whole thalamus was used. The RNA was extracted from KO (*Rosa26*^*CreERt/+*^, *Shox2*^*f/-*^ mice) and CR (*Rosa26*^*CreERt/+*^, *Shox*^*/f/+*^) mice, and RT-qPCR was performed. The Cav3 family of Ca^2+^ channel subunits encode I_T_ and is highly expressed in the nuclei of the thalamus ^56, 57^. We tested the levels of mRNA expression for Cav3.1 and Cav3.2, which are coded by *Cacna1g* and *Cacna1h*, respectively. Previous studies showed that Cav3.1 protein subunits are the primary T-type calcium channel proteins expressed in the thalamocortical neurons, while Cav3.2 proteins are expressed at lower levels in the thalamus and the prominent subunit in the reticular nucleus of the thalamus ^58-61^. Our results showed that expression of *Cacna1h* expression was very low in the thalamus, which confirmed the specificity of our thalamic dissection (Supplemental Fig. 2C) and there was no significant difference in *Cacna1h* expression between CR and KO (Student’s t-test, t_9_=1.02, P=0.34). With respect to the expression of *Cacna1g, Shox2* KO significantly decreased the mRNA expression of *Cacna1g* (Fig. 3C; Student’s t-test, t_9_=3.85, P<0.01) in the thalamic tissue. These results confirmed the mRNA sequencing data and suggest Cav3.1 channels that contribute to the T-type currents are down-regulated in the thalamus.

**Figure 3.**
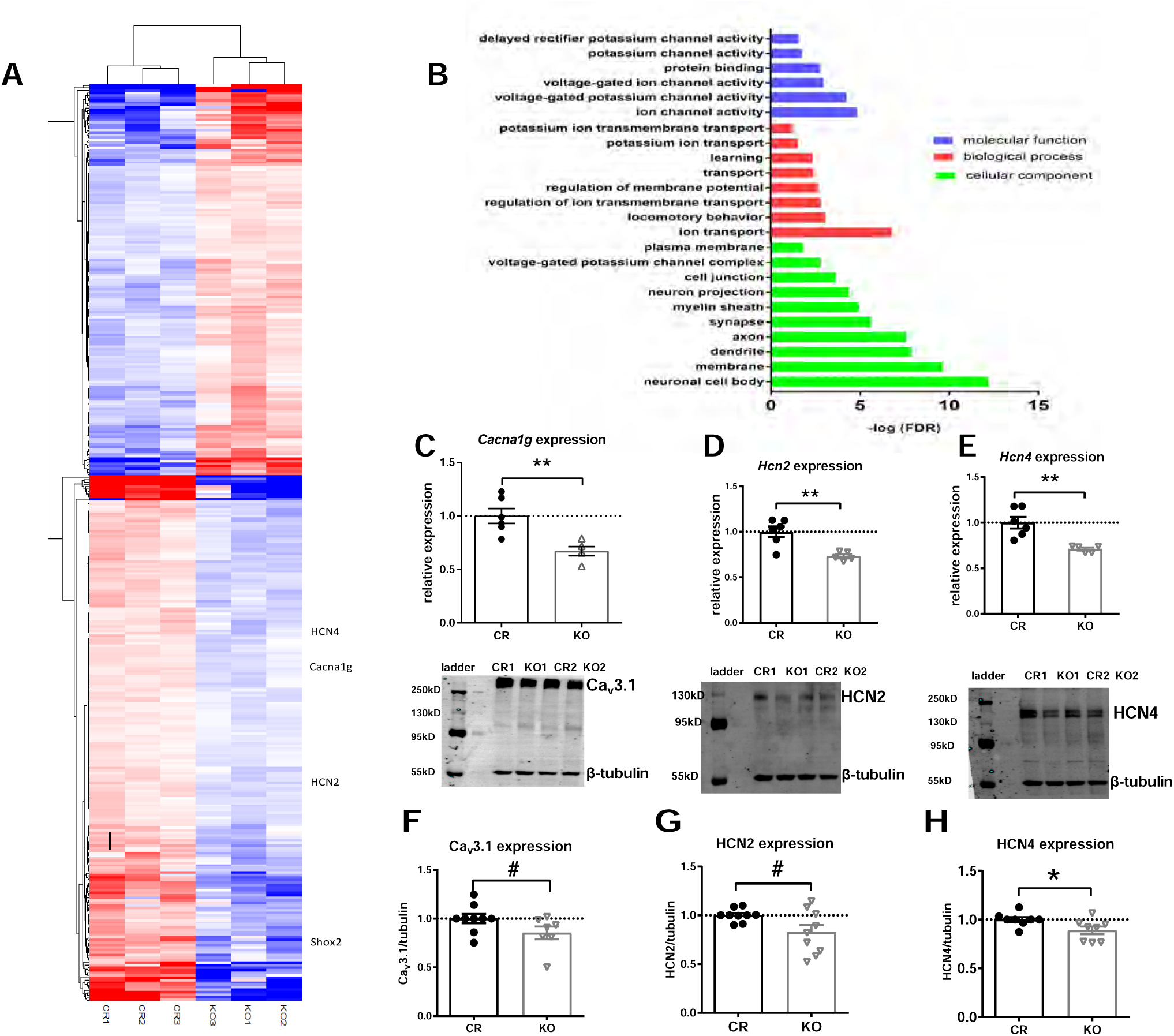
Shox2 expression affects gene expression and ion channel protein levels. RNA-sequencing and analysis were performed as described in methods. **A**. Heatmap, made by pheatmap, saturated at 1, displays 367 DEGs (adjusted p value <0.1) in the medial thalamus between control (CR) and *Gbx2*^*CreErt*,^ *Shox2* KO mice. **B**. Gene ontology (GO) enrichment analysis of DEGs. All terms with an FDR (analyzed by DAVID functional annotation tool) less than 0.1 are listed. **C-E**. RT-qPCR results show that *Shox2* KO significantly reduced mRNA level of *Cacna1g*. **(C)**, *Hcn2* **(D)** and *Hcn4* **(E). F-H**. *Shox2* KO decreased the protein expression levels of Ca_v_3.1 (**F**, ∼120kD), HCN2 (**G**, ∼150kD), and HCN4 (**H**, >250kD) (**, p < 0.01; *, p<0.05, #, p<0.1). The bands around ∼55-60kD are recognized by the β-tubulin antibody.

The mRNA expression of HCN channel genes in CR and KO mice was also assessed. Previous studies reported that mouse brains express very low levels of *Hcn3* ^62^, and our RNA-seq data showed no significant change in *Hcn3* or expression in the KO mice, therefore, *Hcn3* expression was not further investigated. The expression levels of mRNAs for *Hcn1, Hcn2*, and *Hcn4* were investigated. Our results show that *Hcn2* mRNA was the most highly expressed HCN channel gene in the thalamus tissue. *Hcn4* also had prominent expression, while the level of expression of *Hcn1* was only about 5% of *Hcn2* expression. This result is consistent with previous research indicating HCN2 and 4 channels are the most highly expressed HCN channels in the thalamus, and these results along with our sequencing results, provide relative expression data of HCN mRNA expression in mouse thalamus ^62^. *Hcn1* mRNA expression levels were not significantly affected by *Shox2* KO in comparison to CR mice (t_9_=1.85, P=0.10). *Hcn2* and *Hcn4* mRNA were significantly reduced in the *Shox2* KO thalamus compared to CR mice (Fig. 3D, E, *Hcn2*: t_9_=3.92, P<0.01 and *Hcn4*: t_9_=4.02, P<0.01: Mann Whitney nonparametric test). This result also confirmed the RNA-seq results that showed that *Hcn1* mRNA was not significantly affected in mouse thalamus. Together, these results show that *Shox2* KO significantly affects expression of mRNAs for HCN and Ca^2+^ channels. We further investigated if the proteins for these channels were also affected in the KO mice.

Western blot experiments on whole thalamus extract were performed to test the protein levels of the Cav3.1, HCN2 and HCN4 channels. The expression levels of HCN4 proteins were significantly decreased in the thalami of KO animals, and there was a trend toward decreased expression of HCN2 and Cav3.1 proteins compared to CR mice (Fig. 3F,G,H; Cav3.1: t_14_=1.86, P=0.08; HCN2: t_16_=2.30, P=0.1; Mann Whitney nonparametric test; HCN4: t_14_=2.37, P=0.03). The protein measurements in these Western blot staining results are consistent with sequencing and RT-qPCR results and confirmed that HCN4 protein expression is modulated by Shox2 in the adult thalamus. While the change in expression of these channels is relatively small, it’s important to note that these data are taken from the entire thalamus, including neurons and glial cells that do not express *Shox2*. The consistency of the sequencing, mRNA and protein expression is solid evidence that *Shox2* affects expression of these ion channel genes.

Since the expression levels of the channel proteins that underlie currents important for the bursting properties of the thalamic neurons are regulated by *Shox2* expression, we assessed the firing and intrinsic properties of thalamic neurons in *Shox2* KO and CR mice. To best identify a single thalamic nucleus and a homogenous neuron group, the anterior paraventricular thalamus (PVA), the most rostral and dorsal midline nucleus, was chosen as the target region for recording. First, cell-attached voltage-clamp recordings were performed to record the spontaneous action potential currents of PVA neurons without rupturing the cell membrane. Our results indicated that a smaller percentage of cells fired spontaneous action potentials (active neurons) in KO mice compared to those in CR mice (Fig. 4A, B; CR: 36% (9 of 25 cells; N = 3) vs KO: 14% (5 of 35 cells; N = 4; χ2 test, χ2=3.84, P=0.05). Whole-cell patch clamp recordings revealed the decreased cell excitability in PVA neurons from the KO mice was not due to significant differences in resting membrane potential between CR and KO cells (CR: −55.5±1.7 (n=35; N=9) and KO: −54.5±1.7 (n=35; N=10), p = 0.9); however, input resistance was significantly different between cells recorded from KO and CR mice (CR: 993.4 ± 52.9 (n=30; N=9) KO: 850.3±40.7 (n=29; N = 9); p = 0.04). In addition, an increased action potential threshold in KO compared to CR mice was observed (Fig. 4C; t_42_ =2.0; P < 0.05). These results suggest that reduced Shox2 expression affects the firing properties of TCNs.

**Figure 4.**
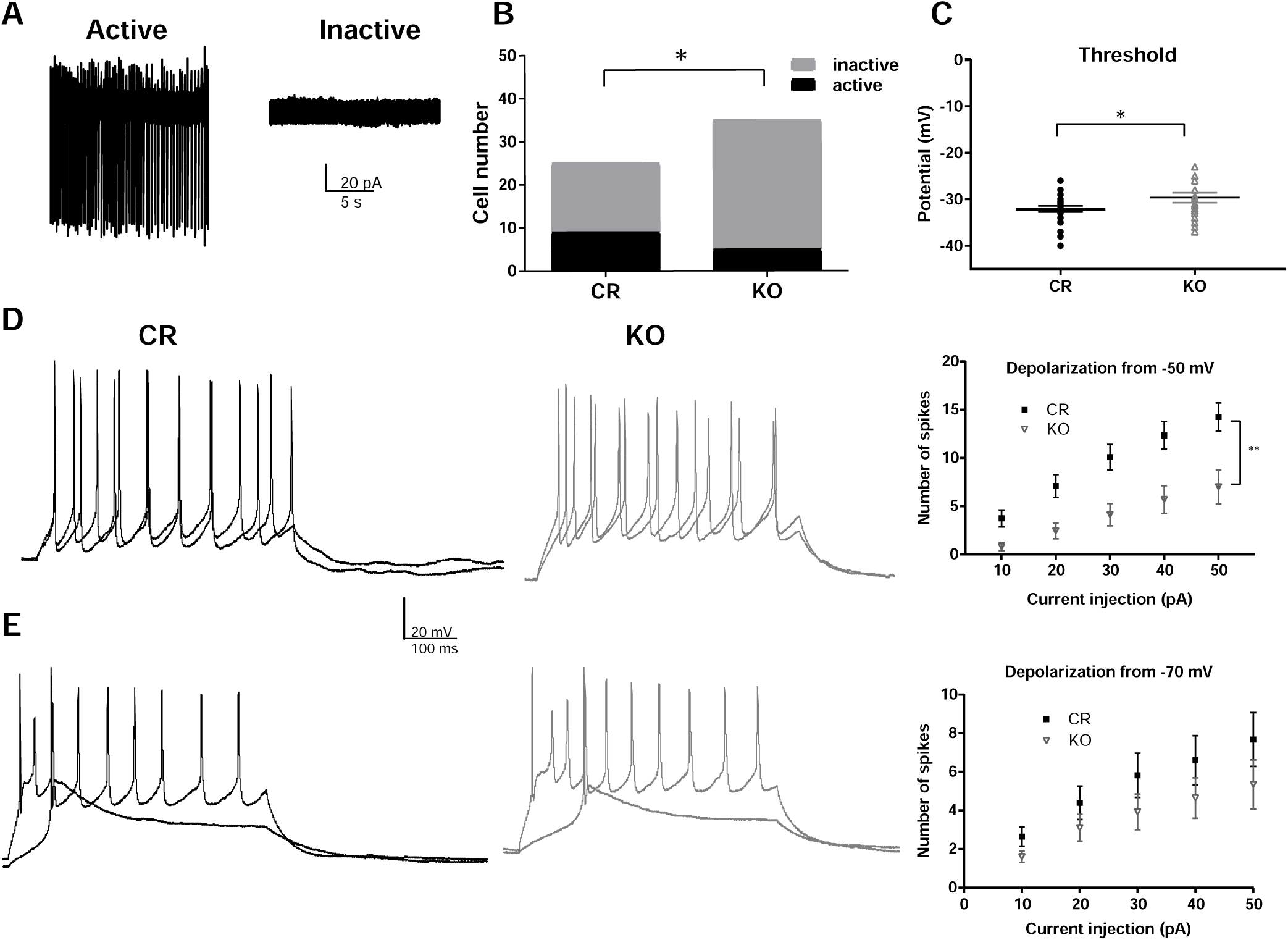
Shox2 KO decreases excitability of TCNs in anterior paraventricular thalamus (PVA). **A**. Example traces of attached-cell recordings of active cells showing spontaneous action potentials (left) and inactive cells with no action potentials (right). **B**. Bar graph representing the ratio of active and inactive cells recorded in PVA from KO and CR mice. This ratio is significantly smaller in KO than in CR mice (*, p<0.05; n = 9 of 25 CR cells (N = 3) and 5 of 25 KO cells (N = 4). **C**. Whole-cell patch clamp recordings showed that action potential threshold was significantly increased in Shox2 KO cells compared to CR cells (KO: n= 16; N= 6; CR: n =23, N= 9; p =0.04). **D**. Example traces of firing patterns triggered by the injection of ramp current (10-50 pA) from near membrane potential (−55 mV) in cells recorded from CR – left, black) and KO mice (right, gray). Right –graph showing reduced spike number in KO cells compared to CR cells at all membrane potentials when depolarized from −55 mV. **E**. Example traces of spike firing triggered by the injection of ramp current (10-50 pA) from −70 mV in cells recorded from CR – left, black) and KO mice (right, gray). Right –graph showing no significant change in spike number in KO cells compared to CR cells at all membrane potentials when depolarized from −70 mV. (*, p < 0.05, **; p < 0.01, ***; p <0.001).

To investigate the spiking responses to depolarizing current injections, we recorded in current clamp mode and injected 10-50 pA currents to evoke action potentials from resting potential (−50-55mV) and −70mV. The number of action potentials fired in response to current injection in neurons recorded from *Shox2* KO slices was significantly decreased compared to that in neurons from CR slices at resting potentials (Fig. 4D; two-way repeat measures ANOVA: main effect of current injection, F_4,148_ =59.9, P<0.0001; main effect of genotype, F_1,37_ =10.9, P<0.01; interaction, F_4,148_ =4.06, P<0.01). Further investigation to determine effects on bursting properties showed that the firing properties in response to depolarizing pulses from −70 mV were not affected in the Shox2 KO cells (Fig. 4E, Two way repeated measures ANOVA: significant main effect of current injection F_4, 216_ = 20.64, P<0.0001; but no significant main effect of genotype CR vs KO, F_1, 54_ = 4.2, P=0.18 or interaction F_4,216_ = 0. 49; P= 0.79). These results are consistent with the spontaneous firing results that suggest reduced *Shox2* affects tonic TCN firing properties at depolarized membrane potentials.

### Shox2 is critical for HCN currents in the thalamus

Since HCN currents play a role in the firing properties of thalamocortical neurons ^74, 75^, and *Shox2* affects expression of *Hcn4* mRNA and protein, we investigated the effect of Shox2 KO on HCN current by sequential hyperpolarizations in voltage-clamp mode in the presence of BaCl_2_ to block inward rectifier K+ currents. The amplitude of HCN current was measured as the difference between the end current of one-second hyperpolarization and the beginning instantaneous current at −150mV hyperpolarization (Fig. 5A). The HCN current densities in neurons from *Shox2* KO mice were significantly decreased compared to neurons from CR mice (t_15_=3.1; P = 0.007; Fig. 5B). Recordings were completed in current clamp to determine the functional impact of reduced I_h_, within the TCNs. Negative current injections from resting potential (Fig. 5C; 10-90 pA in 10 pA steps) induced significantly smaller voltage sags in the KO mice compared to CR mice (Fig. 5C,D; Two way ANOVA; main effect of current input F_(5, 55)_ = 24.5; P<0.0001 and main effect of genotype F _(1, 11)_ = 17.26 but no interaction F _(5, 55)_ = 0.4496; P=0.8). Upon release from the hyperpolarizing pulses, fewer KO cells (2/11) exhibited rebound low threshold Ca^2+^ bursts compared to CR (7/10). These results suggest that Shox2 KO reduced I_h_ and physiologically impacts rebound firing of TCNs, that likely also involves T-type calcium spikes. Significantly, the differences in sag were revealed in the presence of BaCl_2_ and not in control aCSF, which suggests that an inwardly rectifying K^+^ current may partially compensate in the KO for the differences in I_h_.

### Shox2 expression affects TCN synaptic activity

In order to determine the impact of Shox2 KO on synaptic activity, excitatory postsynaptic currents (EPSCs) were measured in cells from CR and KO mice. EPSC interevent interval was significantly decreased in Shox2 KO neurons compared to CR neurons (Fig. 6A-C; t_50_=2.3; P = 0.03), showing increased EPSC frequency in KO slices. Closer investigation revealed instantaneous EPSC frequency was significantly increased in Shox2 KO mice (t_50_ = 3.0; P = 0.004), suggesting increased EPSC burst frequency in slices from KO mice. Cumulative frequency plots showed a significant shift toward increased instantaneous frequency (Kolmogorov-Smirnov, P<0.0001). There was no significant effect on mEPSC amplitude, although a trend was observed (t_50_ = 1.63; P =0.1). These results of increased instantaneous frequency suggest increased burst glutamatergic input to PVA nucleus neurons in the *Shox2* KO mice.

**Figure 5.** Shox2 KO decreased HCN current in anterior PVT of neurons. **A.** An example of HCN current elicited by hyperpolarizing cell membrane from −50mV to −100mV and −150mV. HCN current is defined as the current difference between the current at the end of 1s hyperpolarization and the current peak at the beginning of hyperpolarization as shown in the figure. **B**. Scatter plot showing that *Shox2* KO (n = 9; N = 3) decreased HCN current density in anterior PVT of neurons (**, P<0.01) compared to CR neurons (n = 8; N = 2). **C**. Example current clamp recordings demonstrating inhibitory pulses and sag in CR and KO mice. **D**. Current voltage plot showing sag amplitude measured in response to negative current injection (90-10 pA). (**, P < 0.01; n = 7; N= 2; KO n = 6, N = 3)

**Figure 6.**
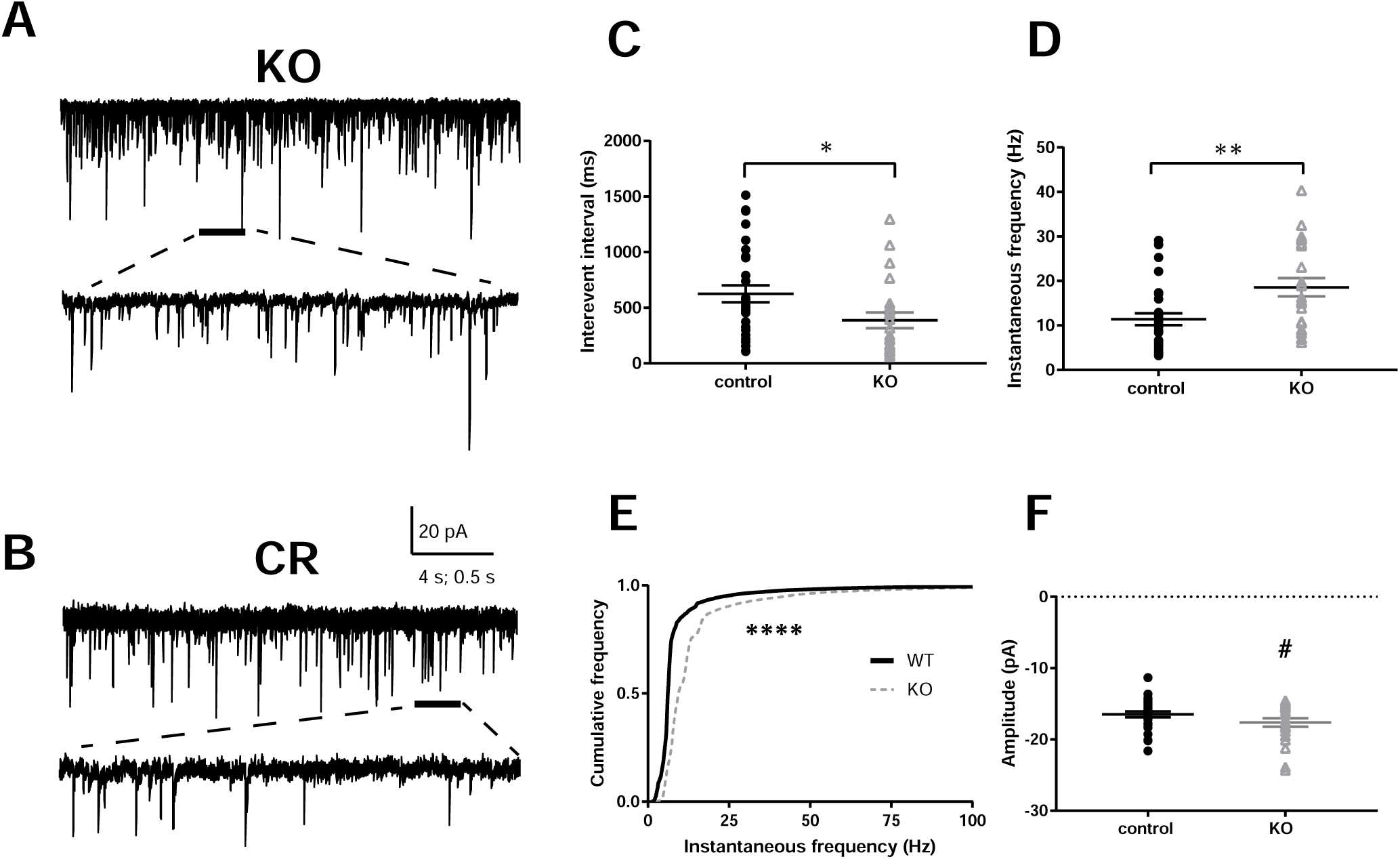
Cells from Shox2 KO mice receive increased frequency of EPSCs. **A**. Example of EPSCs and EPSPs (below) recorded from KO (upper) and CR (lower) mice. **B**. Quantification of interevent interval, Shox2 KO cells showed reduced interevent interval (*, P < 0.05; KO: n = 23; N= 9; CR: n = 29, N = 8). **C**. Quantification of instantaneous frequency, Shox2 KO cells showed increased instantaneous frequency (**, P < 0.01). **D**. Cumulative frequency of instantaneous frequency shows a significant shift toward higher frequencies in KO slices (****, P < 0.0001; Kolmogorov Smirnov test). **E**. No significant effect of Shox2 KO was observed in on EPSC amplitude.

### Shox2 KO induced thalamus-related behavioral deficits in adult mouse

The thalamus plays a critical role in sensory and motor information relay and processing, sleep and arousal, learning and memory, as well as other cognitive functions. The electrophysiological results indicated that *Shox2* KO impaired thalamic burst-related currents, synaptic activity and intrinsic spiking properties. We hypothesized that *Shox2* is critical for proper cognitive and somatosensory behavioral functions in adult mice. To study the specific contribution of *Shox2* expression in the thalamus to behaviors, including anxiety, depression, somatosensory information processing as well as learning and memory, two inducible KO mouse strategies were employed. For the behavioral studies, we used the *Rosa26*^*CreERt/+*^, *Shox2*^*-/f*^ mice line which is a tamoxifen-inducible global *Shox2* KO, and the *Gbx2*^*CreERt/+*^; *Shox2* ^*-/f*^ mice line which is also tamoxifen-inducible but restricts the KO of Shox2 specifically in the midline of the thalamus^76, 77^ and Supplemental Fig. 2.

The total distance travelled in an open field test was measured to investigate the overall activity and general anxiety levels. Distance travelled in the open field in the global *Shox2* KO mice was not statistically significant compared to CR mice (Fig. 7A, t_26_=1.48; p = 0.15). Interestingly, the global KO mice spent a significantly higher percentage of time in the center of the open field compared to CR mice (Fig. 7B, t_26_=2.2; P=0.04), which is indicative of lower levels of anxiety in the KO mice ^78^.

**Figure 7.**
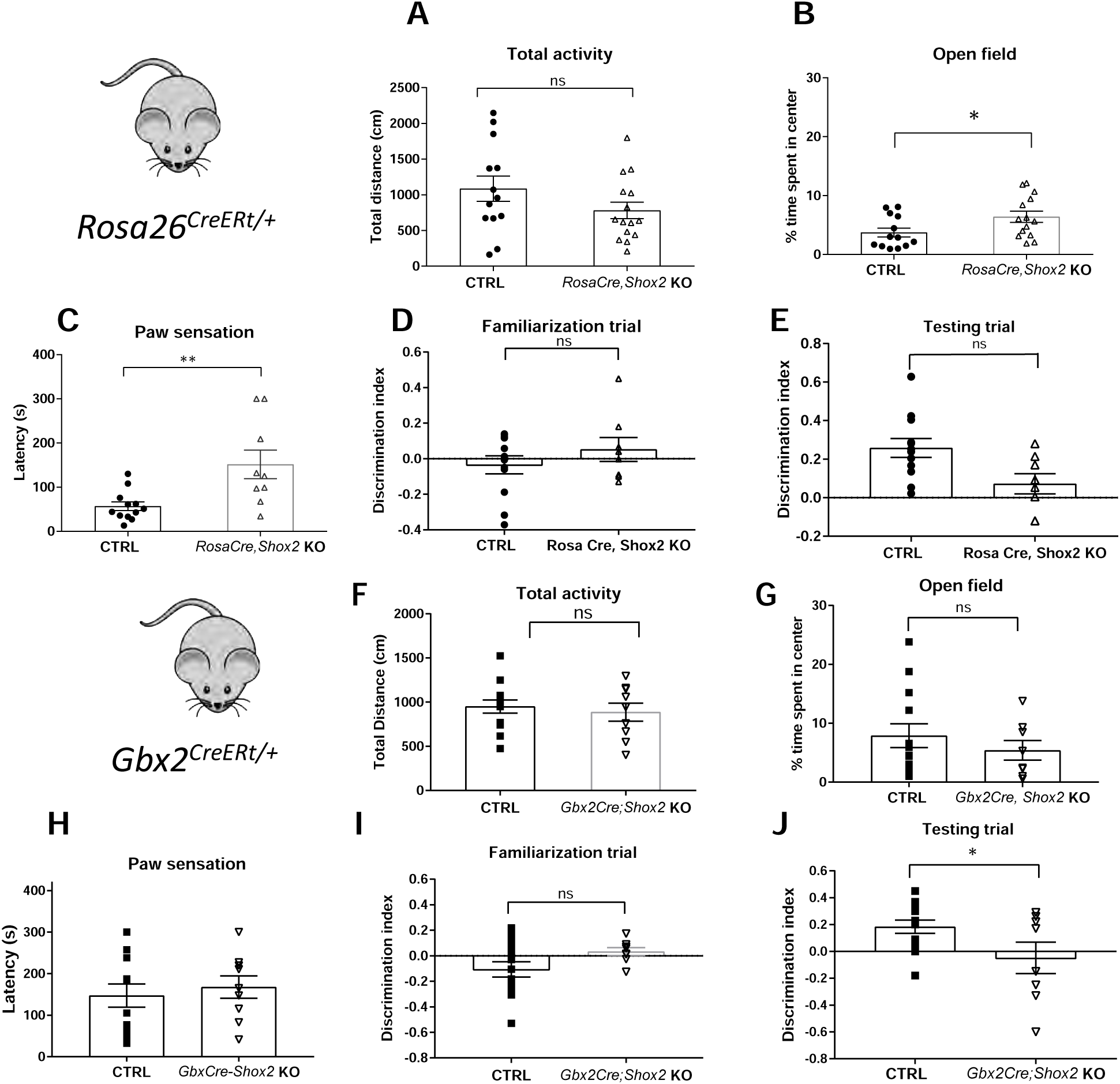
Shox2 inducible KO caused comprehensive thalamus-related behavior deficits. **A**. Results from open field analysis. The total distances travelled by *Rosa*^*CreErt*^, *Shox2* KO and CR mice in 5-minute open field test were similar. **B**. *Rosa*^*CreErt*,^ *Shox2* KO mice spent a higher percentage of time in the center of open field than CR mice (*, P<0.05). **C**. Mice with a ∼1 cm^2^ sticky tape on the left hind paw were placed in home cage and the latency for the mice to first react to the tape was measured. *Shox2* KO mice had a longer latency to first react to the tapes than CR mice (P<0.01). **D-E**. The results of discrimination index showed that *Shox2* KO impaired mice ability in recognizing novel object in testing trial (*, P=0.02), while there was no object preference difference between CR and KO mice in familiarization trial (P =0.33). **F**. The total distance travelled by *Gbx2*^*CreErt*^, *Shox2* KO mice in 5-minute open field test was significantly decreased compared to that by CR mice. **G**. *Gbx2*^*CreErt*,^ *Shox2* KO mice spent a similar percentage of time in the center of compared to CR mice. **H**. *Gbx2*^*CreErt*,^ *Shox2* KO mice were not significantly different from CR in the sticky tape test. **I-J**. Object recognition task in *Gbx2*^*CreErt*,^ *Shox2* KO mice. The results of discrimination index showed that *Gbx2*^*CreErt*,^ *Shox2* KO impaired mice ability in recognizing novel object in testing trial (p < 0.05) (K), while there was no object preference different between CR and KO mice in familiarization trial (J).

Because the open field test results suggested that global *Shox2* KO mice exhibited lower anxiety, we investigated depressive-like behaviors using the tail suspension test and the forced swim test in another cohort of animals ^79^. In these tests, the time during which the animals were actively struggling was measured as mobility time. We observed no significant difference in mobility time between control and global KO mice in either test (Forced swim test, t_20_ =0.39, P=0.70; Tail suspension, t_20_=0.85, P=0.40; Supplemental Fig. 5A,B), suggesting *Shox2* KO did not affect depressive-like behaviors.

To test the performance of the mice in general somatosensory function, the paw sensation test was performed. Sticky tape was applied to the plantar surface of the right hind paw of each mouse, and the latency to the mouse’s first reaction to the tape was measured ^49^. The latency of KO mice to react to the tape was significantly longer than that of CR mice (Fig. 7C; t_26_=2.38, P=0.03). The results suggested that *Shox2* KO induced somatosensory deficits in adult mice.

Given that anterior and medial thalamus are critical for learning and memory processes ^80-83^, we tested a subset of the global *Shox2* KO mice in a novel object recognition test which assesses learning and memory functions ^84, 85^. The test consisted of a familiarization trial and a test trial. During the familiarization trial, we measured the time mice spent exploring 2 identical novel objects (see methods) in the open field environment. Animals of both genotypes explored the 2 objects for similar amounts of time (Fig. 7E, Student’s t-test, t_19_=1.7, P=0.1). Twenty-four hours later, in the memory test trial, the experiment was repeated but one of the beakers was replaced with a new object (Fig. 7D). The percentage of time global *Shox2* KO mice spent around the novel object was significantly decreased compared to that of CR mice in the testing trial (Fig. 7F; t_19_ =2.1, P=0.05). These results suggest an impairment of learning and memory ability of global *Shox2* KO mice.

In order to determine whether the impairment in the object recognition test was mediated by sensory or memory function, we also performed similar behavioral analysis in the Gbx2^CreERt^; *Shox2* KO mice (Fig. 7 G-K), where *Shox2* is reduced specifically in the medial thalamus. The open field test was conducted to investigate the overall activity and general anxiety level of CR and Gbx2^CreERt^; Shox^fl/-^ KO mice. The total distance travelled by *Shox2* KO mice was not significantly different compared to CR mice (Fig. 7G, t_21_=0.92; p = 0.37). In addition, unlike the global KO mice, the time spent in the center of the open field of Gbx2^CreERt^; Shox^fl/-^ KO compared to CR mice was not significantly different (Figure 7H, t_21_=0.50; p = 0.62). We also tested the performance of these mice in general somatosensory function, with the paw sensation test. The latency to react to the tape of KO mice was not significantly different compared to CR mice (Fig. 7C; t_20_=0.4626, P=0.65). The tape fell off the foot of one KO mouse, therefore results from that animal were not used. The results suggested that *Shox2* KO in the medial thalamus did not affect somatosensory function.

We also tested the Gbx2^CreERt^; Shox^fl/-^ mice in the novel object recognition test as described above. Animals of both genotypes explored the 2 objects for similar amounts of time in the familiarization trial (Fig. 7J, Student’s t-test, t_18_=1.64; P=0.11). Twenty-four hours later, in the memory test trial, the experiment was repeated but one of the beakers was replaced with a new object (Fig. 7D). The percentage of time Gbx2^CreERt^; Shox^fl/-^ mice spent around the novel object was significantly decreased compared to that of CR mice in the testing trial (Fig. 7F; t_18_=2.28; P=0.04). These results suggest an impairment of memory formation in the Shox2 KO mice in the medial thalamus, consistent with studies that show the medial thalamus is important for cognitive function.

Since the anterior and medial thalamus have also been implicated in fear memory formation ^86^, cued and contextual fear memory was assessed. Neither contextual (t_21_=0.52; p = 0.61 nor cued fear memory (t_20_=0.1.4; p = 0.17, freezing in one mouse was excluded as an outlier) was affected in the Gbx2^CreERt^; Shox^fl/-^ KO mice (Supplemental Fig. 5C,D). This result is supported by our observations made in td-Tomato animals that show sparse direct inputs to the hippocampus and the amygdala (Fig. 2M-O). Together, these results suggest that the groups of neurons expressing *Shox2* in medial thalamus support recognition memory but are not implicated in fear memory formation or somatosensory information processing.

## Discussion

This study demonstrates the importance of transcriptional activity of the homeobox protein transcription factor, Shox2, in regulation of firing properties, synaptic connectivity, and function of thalamocortical neurons in adult thalamus. This assertion is supported by our investigations at genetic, electrophysiological and behavioral levels. Genetic analysis via RNA-sequencing and Gene Ontology (GO) analysis revealed that Shox2 modulates expression of genes that encode for proteins directly associated with firing properties of TCNs, specifically voltage-gated ion channels. Further investigation using quantitative PCR and Western blotting showed that the mRNAs and proteins for several of these ion channels, namely HCN4 is down regulated in the thalamus of the *Shox2* KO. Electrophysiological analysis showed that *Shox2*-regulation of ion channels modulates the intrinsic firing properties in these neurons, reducing neuronal excitability. In addition, the KO neurons received an increased frequency of EPSCs, suggesting possible presynaptic mechanisms to compensate for decreased membrane excitability to maintain TCN function. Finally, behavioral investigation revealed that global *Shox2* KO mice were impaired in an object memory and somatosensory function test, suggesting that *Shox2* is important to maintain normal function of thalamocortical neurons. In order to discern the somatosensory deficit from the object memory function, we further investigated mice with specific KO of Shox2 in the medial thalamus KO (*Gbx2*^*CreRrt*^*-Shox2*), which maintained Shox2 expression in the lateral thalamus, specifically the VB complex important for somatosensory processing. These mice were impaired in the object recognition task and not sensorimotor functions, suggesting that Shox2 expression in the TCNs of the medial thalamus is important for cognitive function. These studies are consistent with previous results that show lesions to the anterior thalamic nuclei can disrupt object recognition memory in animal models^80, 82, 83, 87^.

Previous clinical studies demonstrated that proper thalamic function is critical for memory formation and consolidation. In humans, damage to the thalamic nuclei, especially medial and anterior nuclei, causes severe memory deficits known as diencephalic amnesia ^12-17^. While the neural circuitry of the effects of Shox2 expression on recognition memory are unclear, perhaps these effects occur via effects on TCN connections to retrosplenial cortex as suggested by the anatomical connectivity indicated in our study (Fig. 2 M-O). The retrosplenial cortex has been linked to temporal order of recognition, also known as ‘what and when’ associations^88^. Future studies are necessary to determine the specific projections and functions of the firing properties of the TCNs involved in these functions.

Our investigations showed that Shox2 KO affected tonic firing properties of TCNs. The functions of the burst and tonic firing properties of thalamocortical neurons are still under investigation. Tonic spike firing mode is thought to contribute to reliable information transfer during perceptive states that conveys sensory information to cortex ^89, 90^. Burst firing mode may allow lack of responsiveness to sensory input during sleep and unconsciousness such as during an absence seizure ^91-94^. On the other hand, recent evidence suggests that thalamic bursts can also occur during awake states and convey a high degree of information about sensory stimuli to serve as a ‘wake-up call’ for cognitive attention ^95-100^. Computational studies suggest that the bursting behavior occurs in response to low-frequency stable inputs, while single spikes occur in response to higher frequency more dynamic input ^101, 102^. Disruptions in the transitions of firing patterns through effects on intrinsic currents in TCNs would disrupt normal thalamic function and its contribution to information processing. Our present studies from the thalamus, together with studies of Shox2 function from the heart ^103, 104^ and excitatory interneurons in spinal cord^34,36^, suggest that *Shox2* is important for maintenance of tonic firing properties in TCNs.

Several lines of evidence indicate that these studies of the role of *Shox2* in pacemaker function in mice are also applicable to humans. *Shox2* is a super-conserved gene with 99% amino acid identity between human SHOX2 and mouse *Shox2*. A recent study found that two missense mutations within the human *SHOX2* gene are associated with early-onset atrial fibrillation, likely caused by a defect in pacemaker activity^105,106^. In addition, while mice do not express the *Shox* gene, human *SHOX* and *SHOX2* have 79% similar amino acid identity, and the same DNA-binding domains and putative phosphorylation sites. The functional redundancy in the regulation of heart pacemaker cells’ differentiation between human *SHOX* and mouse *Shox2* has been demonstrated in mouse models 104, 107. Therefore, investigation of *Shox2* function in mouse can extend to evaluate the role of human *SHOX* and *SHOX2* in humans. Turner syndrome (TS) is one of the most common sex chromosome abnormalities ^108, 109^ and results from the complete or partial loss of the X chromosome. Most individuals with TS have short stature, which is associated with the loss of the SHOX gene ^110-112^. These individuals are at increased risk for neurodevelopmental issues, including learning disabilities, visuo-spatial, social and executive function impairments ^113^ and epilepsy ^114-118^. Interestingly, the smallest chromosomal deletion associated with the neurocognitive phenotype included *SHOX* ^119^, suggesting that loss of SHOX may play a role in cognitive impairments in humans. While the mechanisms of the neurodevelopmental issues in these patients is unclear, our current study indicates that altering expression of SHOX- or SHOX2-related genes may contribute to thalamic dysfunctions and some of these neurodevelopmental impairments.

Further studies are necessary to determine the specific contribution of Shox2-expressing neurons to thalamocortical circuitry, and the role Shox2 may play beyond regulation of firing properties. In addition, future studies will investigate whether *Shox2* plays a critical role during thalamus development and differentiation, the contribution of these *Shox2*-regulated currents to overall thalamocortical neuron function, and the mechanisms by which *Shox2* regulates their expression.

## Supplemental Figures

**Supplemental Figure 1**. Schematic diagrams of breeding schemes used to produce Rosa26^CreERt/CreERt^, Shox2^f/-^ and *Gbx2*^*CreErt/CreErt*^, *Shox*^*f/-*^ mice.

**Supplemental Figure 2. Characterization of *Gbx2* expression and *Gbx2*-induced *Shox2* KO in thalamus. A**. GFP staining (red color) represented the expression pattern of *Gbx2* in P25 mouse brain, showing *Gbx2* only expressed in the midline of the thalamus (arrow shows PVT in midline thalamus) but not lateral thalamus. Blue: DAPI. **B**. qPCR results demonstrating significant knockdown of *Shox2* mRNA in medial thalamus of *Gbx2*^*CreErt/CreErt*^, *Shox*^*f/-*^ mice compared to CR mice (t_6_=3.9, P = 0.008), but no effect on *Shox2* mRNA expression in the lateral thalamus compared to CR animals (**, p<0.01).

**Supplemental Figure 3. Shox2 expression is restricted to thalamus in adult and diencephalon throughout development**. Brain sections demonstrating X-gal staining (or Shox2 expression) results from PND25 **(A1-A8)** and PND56 Shox2^LacZ/+^ **(B1-B4)** and PND56 male Shox2^cre/+^, Rosa26^LacZ/+^ mouse **(C1-C4)**. X-gal staining was observed in anterior thalamus nuclei (ATN), anterior paraventricular nucleus (PVA), ventrobasal thalamus (VB), dorsal lateral geniculate nucleus (dLGN) and medial geniculate nucleus (MGN) but was not observed in the cortex (CX), striatum (STR), hippocampus (HP), amygdala (Ag) or hypothalamus (HT). During development, *Shox2* did express in in habenula (HB), and some areas of the midbrain including superior colliculus (SC) and inferior colliculus (IC), but this expression is reduced in adults. Scale bar: 2 mm.

**Supplemental Figure 4. Strong GFAP immunoreactivity in the hippocampus, but a lack of GFAP+ staining in the thalamus**. Coronal brain section showing strong GFAP+ staining in the hippocampus, but a dearth of GFAP staining in the thalamus.

**Supplemental Figure 5. Behavioral analysis of Shox2 KO mice**. Mobility results measured from CR and *RosaCre;Shox2* KO mice. Forced swim test **(A)** and tail suspension test **(B)**. No significant differences in struggling time between CR and KO mice were observed. **C**,**D**. Freezing behavior measured in cued and contextual fear conditioning in *Gbx2Cre; Shox2KO* mice. No significant differences in freezing behavior to the context or cue were observed.

## Acknowledgements

Funding NIH grants R21NS101482 to LAS and R01 HL136326 to YPC.

## Authorship statement

DY and MM conceived experimental design, performed experiments, and wrote the manuscript. YS, XH, IF, CN, EM, SR contributed data. CS, WY (posthumous) contributed to early planning stages. YPC provided animals and reagents and LAS contributed to overall design and wrote manuscript.

