## Supplementary figures and images for "The transcription factor Shox2 shapes thalamocortical neuron firing and synaptic properties"

### supplemental figures

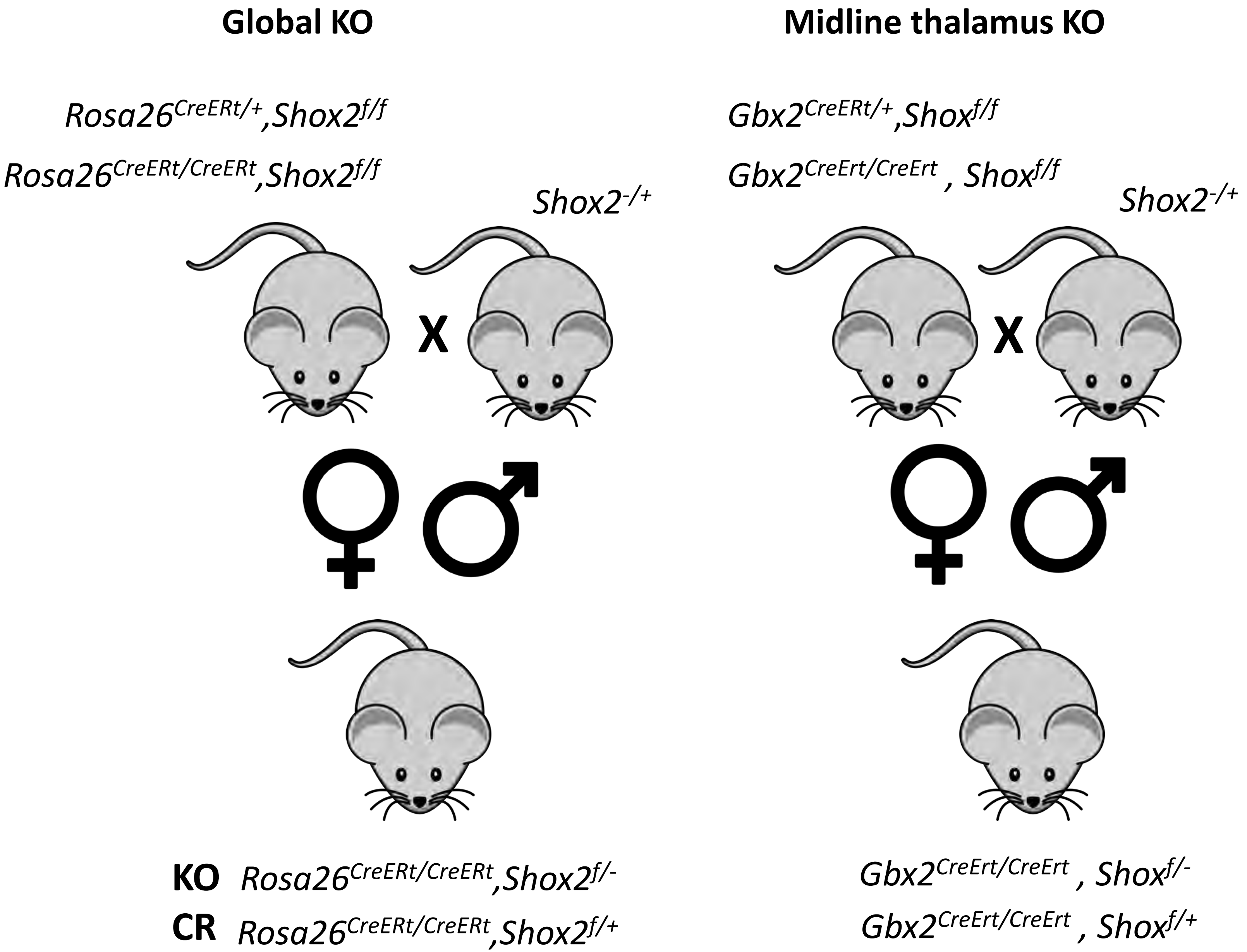

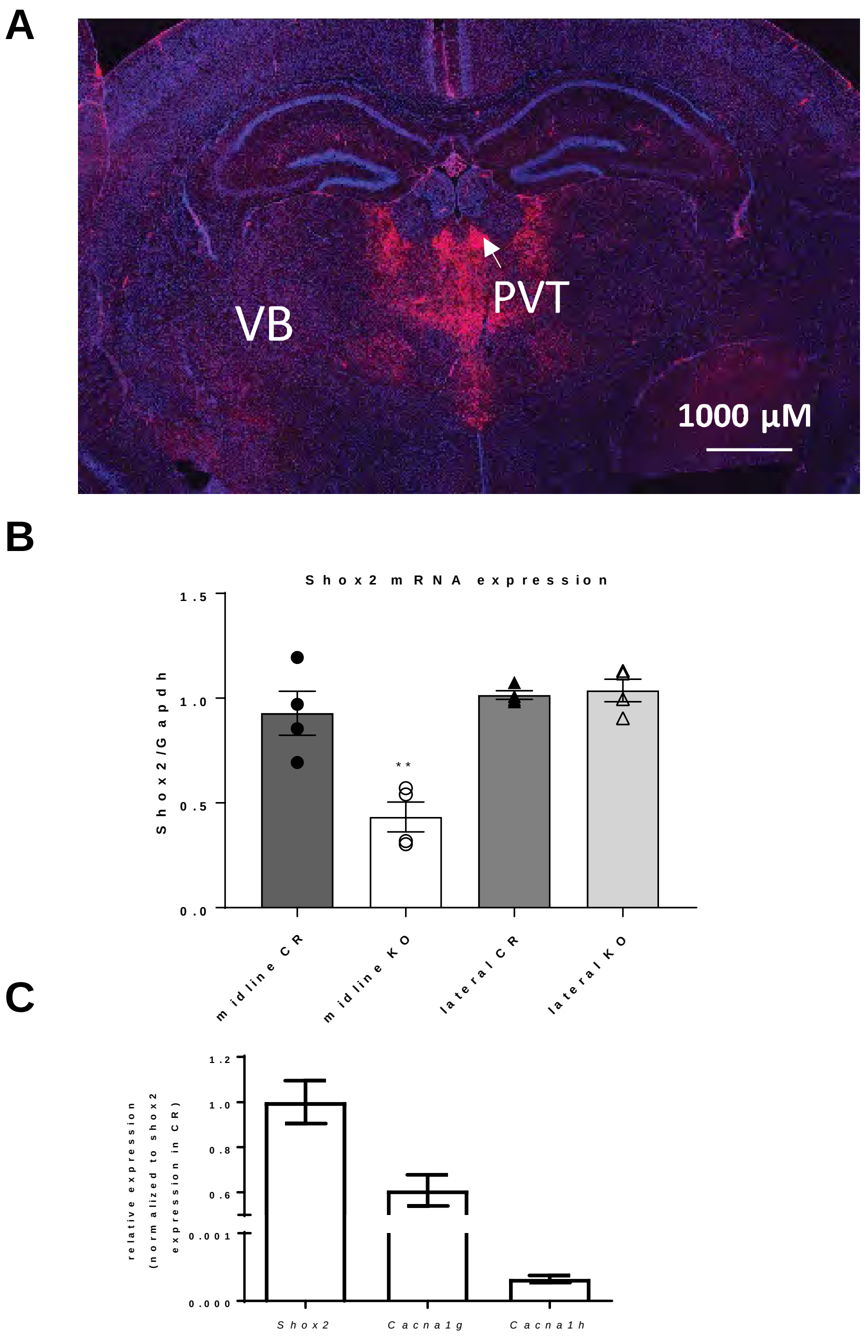

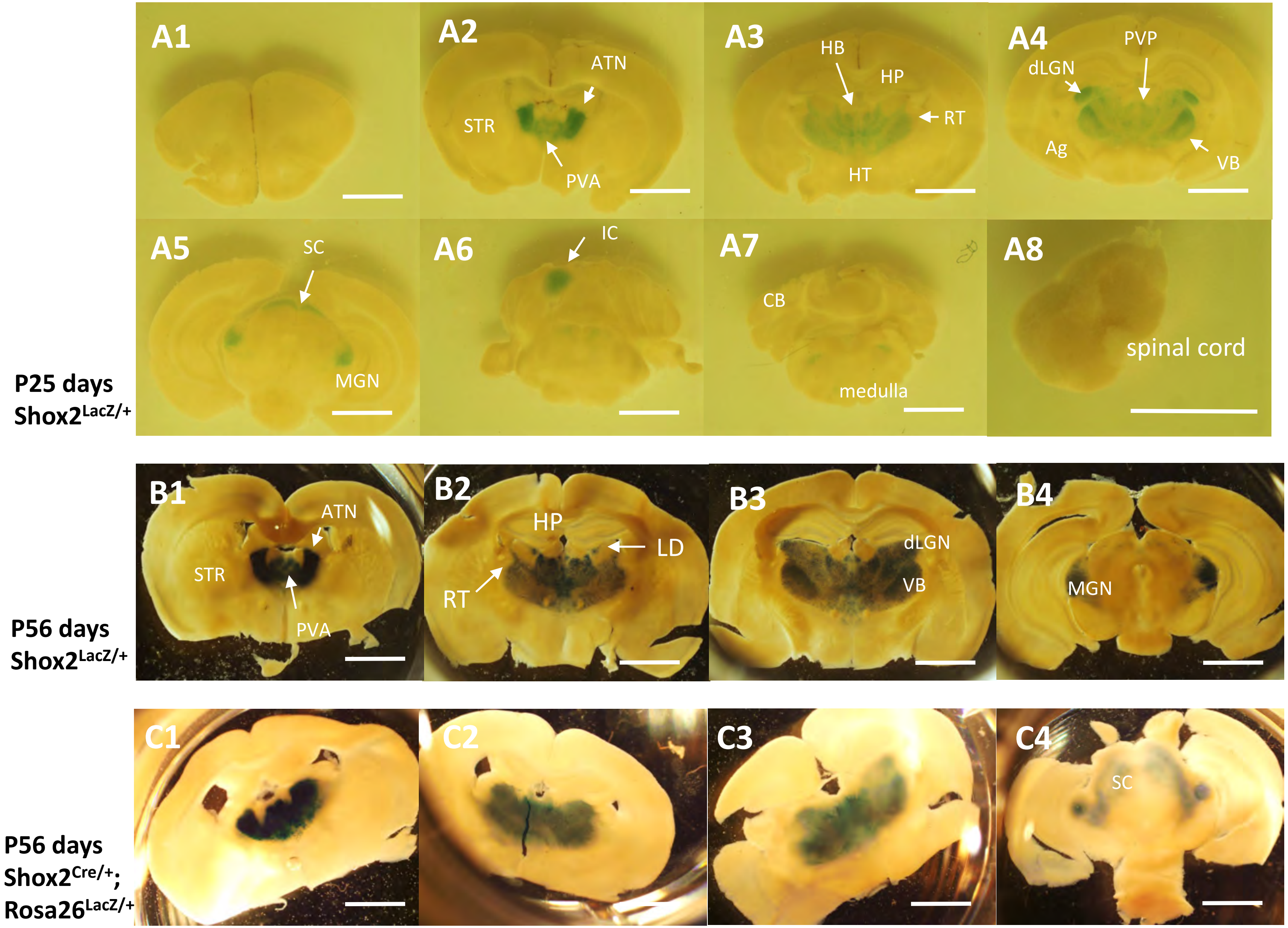

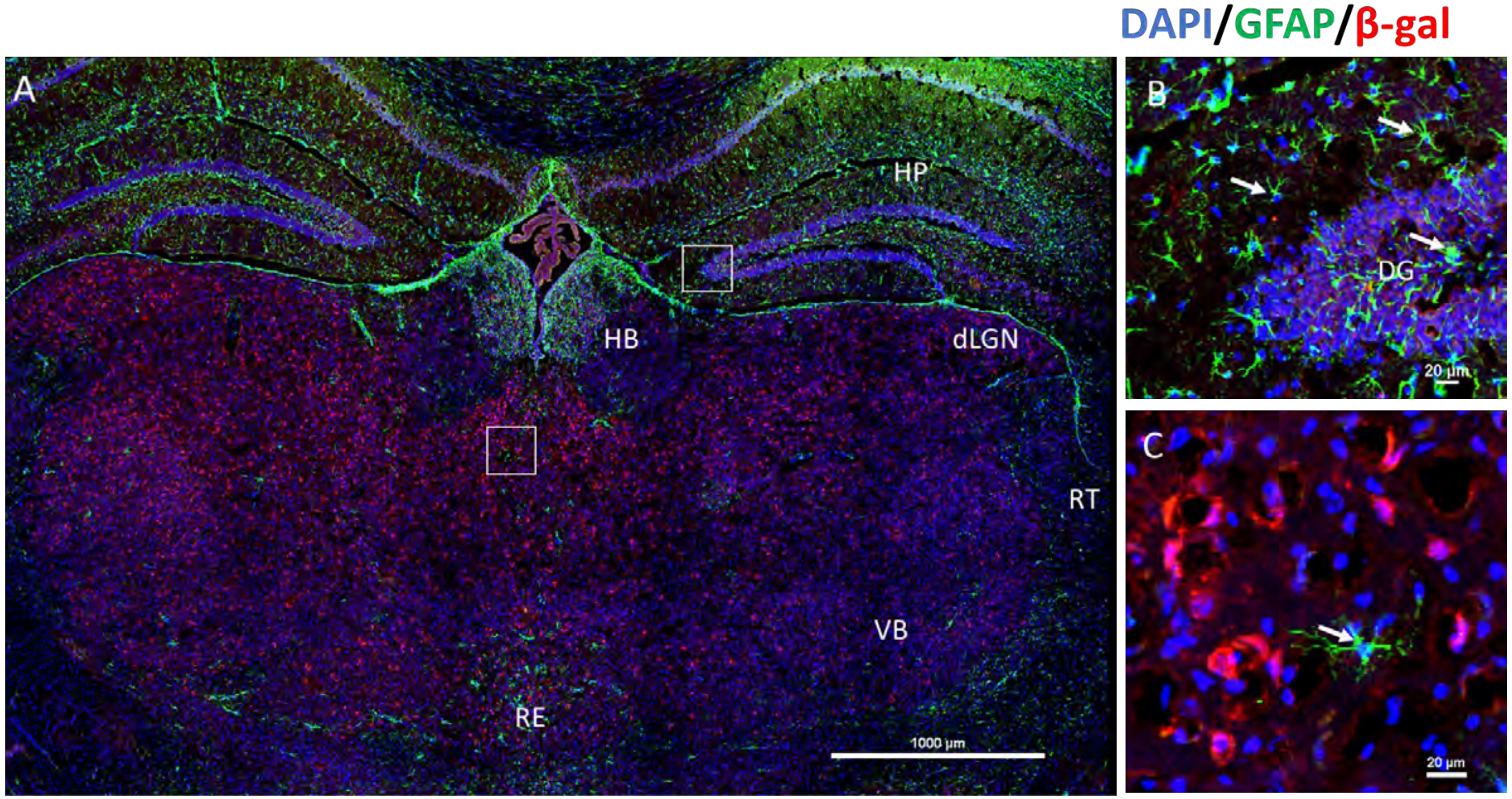

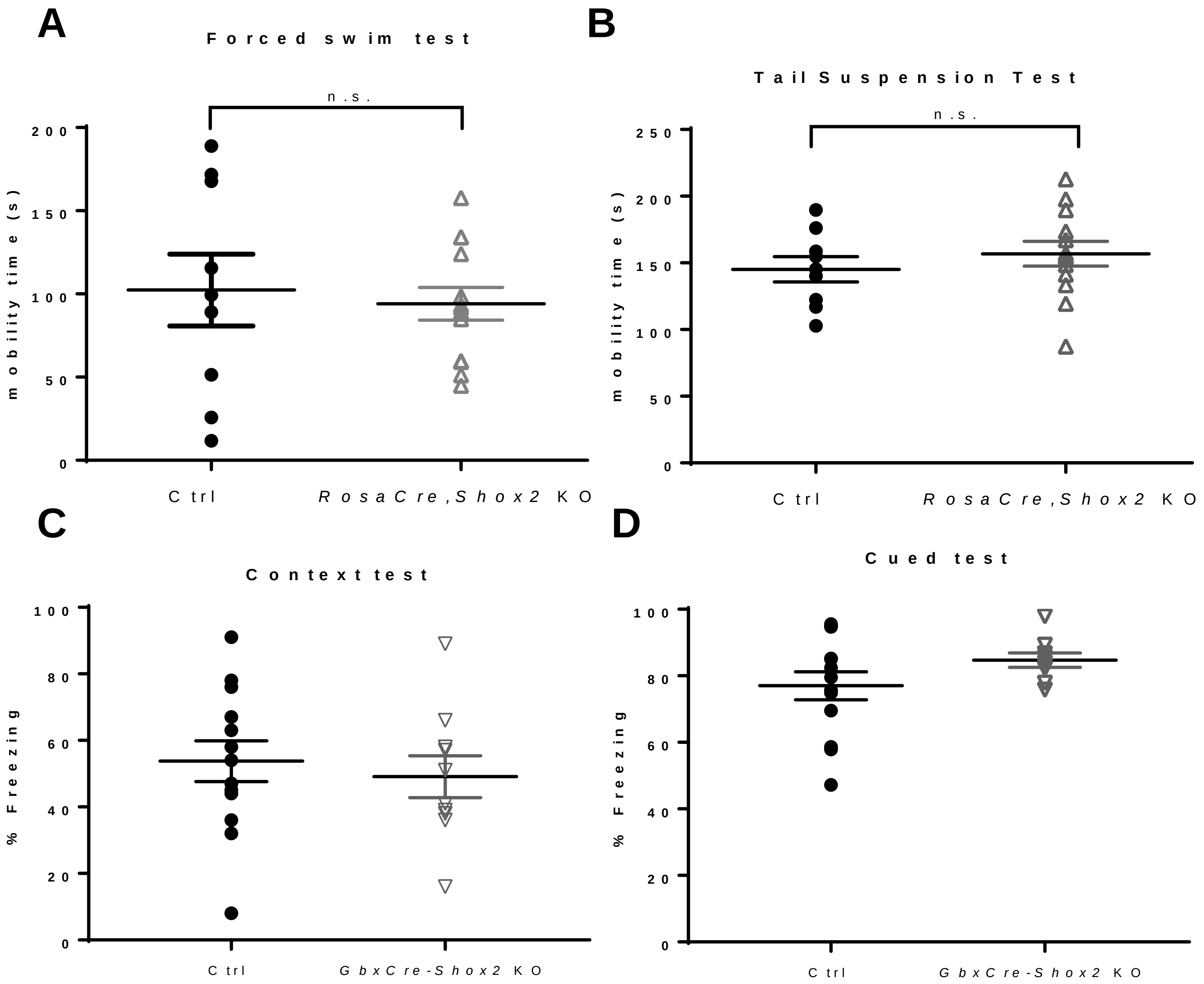
